# Natural variation suggests candidate genes underlying *Caenorhabditis elegans* susceptibility to diverse toxicants

**DOI:** 10.1101/2025.11.30.691394

**Authors:** Timothy A. Crombie, Ryan McKeown, Samuel J. Widmayer, Amanda O. Shaver, Nicolas D. Moya, J. B. Collins, Janneke Wit, Robyn E. Tanny, Christian Braendle, Lewis Stevens, Lisa van Sluijs, Matthew V. Rockman, Mark G. Sterken, Marie-Anne Félix, Erik C. Andersen

**Affiliations:** Department of Biomedical Engineering and Science, Florida Institute of Technology, Melbourne, FL; Molecular Biosciences, Northwestern University, Evanston, IL; Interdisciplinary Biological Sciences Program, Northwestern University, Evanston, IL; Department of Biology, Johns Hopkins University, Baltimore, MD; Program in Cell, Molecular, Developmental Biology, and Biophysics, Johns Hopkins University, Baltimore, MD; Université Côte d’Azur, CNRS, Inserm, IBV, Nice 06100, France; Wellcome Sanger Institute, Tree of Life, Wellcome Genome Campus, Cambridge, CB10 1SA, United Kingdom; Laboratory of Nematology, Wageningen University & Research, Wageningen, the Netherlands; Department of Biology and Center for Genomics & Systems Biology, New York University, New York, NY; Institut de Biologie de l’École Normale Supérieure, CNRS, Paris, France

**Keywords:** *Caenorhabditis elegans*, genome-wide association studies (GWAS), environmental toxicants, xenobiotic response, risk assessment

## Abstract

Genetic differences among individuals shape how they respond to environmental toxicants, but the identification and validation of the genes responsible for this variation is difficult, particularly in humans. Consequently, our limited knowledge of the genes that influence susceptibility constrains our ability to accurately predict the risks posed by environmental toxicants. To identify genes underlying natural differences in toxicant susceptibilities, we measured the effects of 23 environmental toxicants on larval development across 195 genetically diverse *Caenorhabditis elegans* strains using a high-throughput imaging platform. We then combined these response data with whole-genome sequences to perform genome-wide association mappings, identifying 40 genomic regions where genetic variants are correlated with susceptibility differences. Many of these regions are enriched for genes involved in biological processes previously linked with toxicant responses, supporting the potential contributions of these genes to natural variation in susceptibility. Using a set of heuristics, we identified 94 candidate susceptibility genes, offering targets for experimental validation that could ultimately inform toxicant risk prediction and regulatory assessment by linking genetic variation to differences in susceptibility.

**Impact Statement:** Analysis of natural genetic variation among 195 wild *C. elegans* strains identified 94 candidate genes putatively linked to differences in susceptibility to 23 environmental toxicants. These findings can inform the discovery of conserved susceptibility genes and the development of biomarkers that improve chemical risk assessment by accounting for genetic differences among humans.

## 1 Introduction

It is estimated that more than 350,000 chemicals or chemical mixtures are registered for production and use globally, yet the risks that most of these chemicals pose to human health remain unknown (Wang *et al*., 2020). Accurately characterizing chemical toxicity in humans is challenging, prompting researchers to rely on data from toxicity assays using human cell lines or animal models to predict risk (Rusyn *et al*., 2010; Zeise *et al*., 2013; Chiu and Rusyn, 2018; Rusyn *et al*., 2022). However, most of these assays are performed in a single genetic background, limiting the ability of researchers to observe variability in adverse responses to environmental chemicals found in natural populations (Rusyn *et al*., 2022). Consequently, the full spectrum of genetic and molecular mechanisms that contribute to differences in toxicant susceptibility among genetically diverse individuals remain largely unknown (Zeise *et al*., 2013; Alam and Jones, 2014; Chiu and Rusyn, 2018; Rusyn *et al*., 2022). This critical knowledge gap hinders efforts to understand the genetic determinants of susceptibility, recognize at-risk individuals, and establish evidence-based regulatory limits for chemicals (Zeise *et al*., 2013; Chiu and Rusyn, 2018; Rusyn *et al*., 2022; To *et al*., 2023). A key step in closing this gap is the identification of genes that influence toxicant susceptibility.

Genome-wide association studies (GWAS) are a powerful method to identify genes that contribute to differences in toxicant susceptibilities among individuals. In a GWAS, researchers test for statistical associations between genetic variants, typically single-nucleotide variants (SNVs), and variation in a phenotype, such as response to a toxicant, across a population of genetically diverse individuals (Barton and Keightley, 2002; Broekema *et al*., 2020). These “marker” variants are distributed throughout the genome and are used to scan for genomic regions statistically associated with population-wide differences in susceptibility. In many cases, the associated regions contain variants that impact genes or regulatory elements that can alter protein function, gene expression, or molecular pathways relevant to toxicity. However, because GWAS identifies statistical correlations rather than direct causal relationships, experimental validation is required to determine whether candidate genes in associated regions truly affect toxicant susceptibility.

Human GWAS have identified a limited number of candidate genes associated with toxicant susceptibility by using toxicant levels in blood or urine as the focal trait. The susceptibility genes uncovered typically encode drug-metabolizing enzymes (Pierce *et al*., 2012; Evans *et al*., 2013) or toxicant-interacting proteins such as cellular transporters and metalloenzymes (Moksnes *et al*., 2024; Ng *et al*., 2015; Wahlberg *et al*., 2018). However, identification and validation of candidate susceptibility genes in humans are heavily constrained by confounding variables and ethical restrictions, including age, sex, environmental heterogeneity, and the inability to edit genes in human subjects (Alam and Jones, 2014). To overcome these challenges, genetically diverse human cell lines have been used to assay toxicity, revealing candidate genes involved in solute transport, membrane function, immune response, cell division, and DNA replication (Abdo, Xia, *et al*., 2015; Abdo, Wetmore, *et al*., 2015; Ford *et al*., 2022). Although these *in vitro* models offer valuable insights into the genetic determinants of susceptibility, they are unable to capture the full complexity of whole-organism toxicant responses. For example, cell lines do not express all genes relevant to toxicant responses and the complex interactions between tissues and organ systems are absent (Wheeler and Dolan, 2012). Recent advances in organoid technologies partially overcome these limitations by more accurately modeling the structural and functional complexity of human tissues, but current organoid models still lack the genetic diversity required to study population-level variation in susceptibility (Rossi *et al*., 2018; Co *et al*., 2023). Because of the limitations of human experiments, animal models are indispensable for conducting GWAS aimed at identifying genetic factors that influence toxicant susceptibility at the whole-organism level.

Several genetically diverse animal populations have been used to identify genetic factors that contribute to differences in susceptibility among individuals (Rusyn *et al*., 2022). For example, studies using genetically diverse rodent populations have identified candidate susceptibility genes involved in xenobiotic metabolism (Church *et al*., 2014; Nguyen *et al*., 2017; Yoo *et al*., 2015; You *et al*., 2020), immune response (Harrill *et al*., 2009), chromatin structure (Koturbash *et al*., 2011), and tissue repair pathways (Harrill *et al*., 2012; Tsuchiya *et al*., 2012; Harrill *et al*., 2018). Although genetically diverse rodent populations are a powerful mammalian model for toxicological studies, the model is inherently low-throughput, making it less suited for large-scale experimentation. In addition, high costs and ethical concerns surrounding animal welfare are significant barriers in rodent-based studies (Akhtar, 2015). To address these barriers, the U.S. Environmental Protection Agency (EPA) has outlined a strategic plan to advance new approach methodologies (NAMs), including *in silico*, *in chemico*, and *in vitro* technologies, to reduce reliance on traditional animal models in toxicant risk assessment (USEPA, 2018; Schmeisser *et al*., 2023). Similarly, the National Institutes of Health (NIH) has emphasized NAMs, either alone or in combination with data from tractable animal models, to study disease (NIH to Prioritize Human-Based Research Technologies). In this context, small model organisms provide an attractive non-mammalian alternative, providing higher throughput, reduced cost, and limited ethical concerns that can better keep pace with rapid chemical production. Moreover, genetically diverse populations of *Drosophila melanogaster (Zhou et al., 2017; Everman and Macdonald, 2024; Hanson et al., 2025)*, *Danio rerio (Balik-Meisner et al., 2018; Thunga et al., 2022)*, and *Caenorhabditis elegans* (Vertino *et al*., 2011; Andersen *et al*., 2015; Evans *et al*., 2018; Zdraljevic *et al*., 2019; Bernstein *et al*., 2019; Evans *et al*., 2020), have already been successfully used to screen toxicant responses and identify genetic factors that contribute to inter-individual differences in susceptibility.

*C. elegans* is a powerful whole-animal model for studying toxicant responses that has helped reveal the molecular mechanisms of toxicity for many compounds (Martinez-Finley and Aschner, 2011; Hartman *et al*., 2021). Several features make *C. elegans* a good fit for identifying genes that cause differences in toxicant susceptibility with potential relevance to human health. First, conservation of genes and biological pathways between *C. elegans* and humans is high (Kaletta and Hengartner, 2006), and toxicant responses in *C. elegans* are often predictive of mammalian responses (Boyd *et al*., 2016; Hunt, 2017). Second, *C. elegans* is easily reared in the laboratory, has a short life-cycle (Corsi *et al*., 2015), and several high-throughput phenotyping platforms have been developed that measure a range of toxicity endpoints, including fecundity (Andersen *et al*., 2015), development (Andersen *et al*., 2015; Widmayer, Crombie, *et al*., 2022), and behavior (Boyd *et al*., 2007; Barlow *et al*., 2022). Third, the *Caenorhabditis* Natural Diversity Resource (CaeNDR) provides a public repository of genetically diverse strains, along with whole-genome sequences and genomic tools to help dissect traits, including traits relevant to toxicology (Crombie *et al*., 2024). Fourth, genetically diverse *C. elegans* strains exhibit significant differences in their responses to toxicants and a large proportion of that variation is attributable to genetic differences (Vertino *et al*., 2011; Andersen *et al*., 2015; Evans *et al*., 2018; Bernstein *et al*., 2019; Zdraljevic *et al*., 2019; Widmayer, Crombie, *et al*., 2022; Leuthner *et al*., 2024), enabling statistically powerful GWAS (Widmayer, Evans, *et al*., 2022) to identify multiple genomic regions associated with natural differences in toxicant response (Vertino *et al*., 2011; Andersen *et al*., 2015; Evans *et al*., 2018; Zdraljevic *et al*., 2019; Evans *et al*., 2020; Bernstein *et al*., 2019).

A clear example of the translational potential of *C. elegans* research comes from a recent GWAS that identified natural variation in *dbt-1*, a gene involved in branched-chain amino acid metabolism, as a determinant of arsenic susceptibility (Zdraljevic *et al*., 2019). Subsequent experiments in human embryonic kidney cells (HEK293T) showed that natural variation in the human ortholog of *dbt-1* similarly mediates arsenic-induced changes in fatty acid metabolism, revealing a novel and conserved mechanism of toxicity that may contribute to population-level differences in human arsenic susceptibility (Zdraljevic *et al*., 2019). Here, motivated by the translational potential of *C.elegans*, we sought to uncover candidate susceptibility genes for 23 environmental toxicants with substantial human health implications. The 23 diverse compounds represent five use groups (metals, herbicides, fungicides, insecticides, and flame retardants) and were selected using criteria outlined in the Toxic Substances Control Act, as amended in 2016 by the Frank R. Lautenberg Chemical Safety for the 21st Century Act, which prioritizes compounds with significant human exposure potential, environmental persistence, and bioaccumulation tendencies, among others (US Congress, 2016; McPartland *et al*., 2022). Additionally, we prioritized toxicants for which susceptibility is known to be genetically determined in *C. elegans*, based on previous dose-response experiments (Widmayer, Crombie, *et al*., 2022). To identify candidate susceptibility genes for these 23 environmental toxicants, we performed a GWAS using a cohort of 195 wild strains. Differences in toxicant susceptibilities among the strains were quantified using previously described high-throughput techniques (Widmayer, Crombie, *et al*., 2022; Shaver and Andersen, 2025). In total, we identified 40 genomic regions that correlate variation in thousands of genes with differences in toxicant susceptibilities. Guided by heuristics informed by two decades of quantitative trait research in *C. elegans*, we identified and prioritized 22 actionable genomic regions containing 94 candidate genes that can be experimentally validated as susceptibility genes using the organism’s robust genetic toolkit (Andersen and Rockman, 2022). Future studies that link these genes to toxicant susceptibility could reveal conserved susceptibility genes with translational relevance, ultimately enhancing chemical risk assessment for humans.

## 2 Materials and Methods

### 2.1 *C. elegans* strains

The 195 strains for the mapping panel (**Table S1**) were sourced from CaeNDR and selected based on three criteria (Crombie *et al*., 2024). First, we chose 11 strains (PD1074, CB4856, CX11314, DL238, ECA36, ECA248, ECA396, JU775, MY16, RC301, XZ1516) because they were included in one or both of two recent dose-response assays (Widmayer, Crombie, *et al*., 2022; Shaver *et al*., 2023). Importantly, these dose-response assays used the same phenotyping platform as this study, which allows for direct comparisons of experimental results. In CaeNDR, strains that share alleles at 99.997% of variant sites are grouped into genome-wide haplotypes called isotypes. Of the 11 strains referenced above, two (PD1074 and ECA248) are referred to by their isotype names (N2 and CB4855, respectively) in our analyses (Yoshimura *et al*., 2019; Ichikawa *et al*., 2025). All but three of the remaining 184 strains in the mapping panel were chosen based on the second criterion, assessments of detection power and false discovery rate (FDR), that were made using the NemaScan (*9263c98*) simulation profile, which is an open-source pipeline for GWAS mapping in *C. elegans* (Widmayer, Evans, *et al*., 2022). In general, as the number of strains included in a mapping panel increases, the power of GWAS to detect simulated causal variants increases and the FDR decreases. Moreover, mapping panels with less population structure have substantially reduced FDR (Widmayer, Evans, *et al*., 2022). For these reasons, we chose the largest mapping panel that we could phenotype at high replication (192 strains). To optimize the strain composition of the mapping panel, we generated 50 different hypothetical 192 strain mapping panels by randomly sampling 181 strains to pair with the 11 strains chosen above. We then assessed differences among the 50 mapping panels in detection power and FDR across a range of simulated causal variants. We simulated 30 traits per strain panel, each with narrow-sense heritabilities of 0.2, 0.5, or 0.8. For each trait, variation was attributed to five causal variants with minor allele frequencies (MAF) above 0.05. The effect size of each causal variant was randomly sampled from a gamma distribution. Ultimately, we chose the mapping panel with the best combination of high detection power and low FDR (**Figure S1**). Finally, we included three additional strains (AB1, MY1, PB306) to increase overlap with the *Caenorhabditis elegans* Multiparental Experimental Evolution (CeMEE) panel to better enable future comparisons of GWAS results across strain resources (Noble *et al*., 2017). All nematode strains were maintained at 20°C on 6 cm plates with modified nematode growth medium (NGMA), containing 1% agar and 0.7% agarose to prevent burrowing (Andersen *et al*., 2014). The NGMA plates were seeded with the *Escherichia coli* strain OP50 as a nematode food source. All strains were grown for three generations without starvation on NGMA plates before toxicant exposure to reduce the transgenerational effects of starvation.

### 2.2 Liquid culture bacterial food preparation

Three large batches of HB101 *E. coli* were prepared as a food source for liquid culture of nematodes during toxicant exposures. For each batch of food, a starter culture was seeded with a single bacterial colony and grown in a sterile culture tube with 5 mL of 1X Horvitz Super Broth (HSB) at 37°C with shaking at 180 rpm. After 18 hours, the optical density of the seed culture was measured (OD600) and used to inoculate fourteen 4 L culture flasks at an initial OD600 value of 0.001 in 1 L of 1X HSB. These cultures were then grown at 37°C with shaking at 180 rpm for 15 hours. Afterwards, the culture flasks were moved to 4°C to arrest growth, and the bacterial cells were washed as previously described (Widmayer, Crombie, *et al*., 2022). Finally, the washed bacterial cells for all fourteen culture flasks were resuspended in K medium (51 mM NaCl, 32 mM KCl, 3 mM CaCl2, 3 mM MgSO4, 1.25 μg/mL Cholesterol) (Boyd *et al*., 2009) and combined at a final OD600 value of 100 as a single batch, aliquoted into 15 mL conicals, and frozen at -80°C for later use. To avoid batch effects caused by subtle differences in bacterial growth dynamics, frozen aliquots from each of the three batches were thawed and mixed in equal proportions prior to all feedings, as done previously (Gouveia *et al*., 2021).

### 2.3 Toxicant stock preparation

We prepared stock solutions of all toxicants with water or dimethyl sulfoxide (DMSO), depending on the aqueous solubility of the toxicant. The exact sources, catalog numbers, stock concentrations, and preparation notes for each of the toxicants are provided (**Table S2**). Following preparation of the toxicant stock solutions, they were aliquoted to microcentrifuge tubes and stored at -20°C for use in the high-throughput toxicant response assay. The exposure concentration used for each toxicant in the GWAS was taken from the dose with the highest heritability estimate in a recent dose-response assay (Widmayer, Crombie, *et al*., 2022). We also chose to include a low and high dose for some toxicants in the GWAS (**Table S2**). A secondary dose was included in the GWAS only if the rank order of strain responses deviated between high and low exposure concentrations in the Widmayer *et al*. dose-response assay and narrow-sense heritability estimates were greater than 0.2 for both doses. The logic for including both doses is that distinct genetic mechanisms could underlie the differential strain responses at the two concentrations tested.

### 2.4 Toxicant use groups and mechanisms of action (MoA)

We created toxicant assessment groups based on previously published use categories and mechanisms of action (MoA) data (Kramer *et al*., 2024). First, we queried the ECOTOXicology Knowledgebase (Olker *et al*., 2022) with toxicant CAS numbers to retrieve DSSToX Substance Identifiers (DTXSID), which we used to extract the chemical use and MoA data (Kramer *et al*., 2024). For use-based classification, we assigned the 23 toxicants to five distinct groups: flame retardants (N = 1), fungicides (N = 4), metals (N = 8), herbicides (N = 3), and insecticides (N = 7) (**Table S2**). For three toxicants (mancozeb, paraquat, and silver nitrate), use category data were not available, and use groups were manually assigned based on information from regulatory and manufacturer documentation. For MoA-based classification, we applied the specific categories from Kramer et al. (2024) to group toxicants that act on similar biological targets or pathways. This approach yielded three primary MoA groups: acetylcholinesterase (AChE) inhibition (N = 6), redox balance disruption (N = 10), and other or unknown (N = 7) (**Table S2**). MoA assignments were refined for several compounds based on available literature. Mancozeb was classified as an AChE inhibitor (Melnikov *et al*., 2023). Paraquat, mercury, and cadmium were categorized as redox balance disruptors based on previous studies (Cirovic and Satarug, 2024; Wang *et al*., 2019; Ke *et al*., 2023). Chlorothalonil, originally categorized as “Multi-site activity”, was reclassifiedas a toxicant that affects redox balance based on evidence from marine nematodes and zebrafish (da Silva Barreto *et al*., 2018, 2020).

### 2.5 High-throughput toxicant response assay

We measured toxicant response and control phenotypes for 195 strains across 28 experimental conditions (two control conditions and 26 toxicant conditions) at high replication using a previously described high-throughput, image-based phenotyping assay (Widmayer, Crombie, *et al*., 2022; Shaver *et al*., 2023; Shaver and Andersen, 2025). To accommodate the large number of treatments in our design (195 strains x 28 conditions = 5,460 treatments), we measured toxicant responses over ten sequential experimental blocks. Each block contained all 28 experimental conditions. The first nine blocks contained 20 unique strains each and the tenth block contained 11 unique strains. Four strains (CB4856, JU775, MY16, and N2) were included in all ten blocks to help estimate block effects. For each experimental block, strains were passaged for three generations, then bleach-synchronized in triplicate to account for any variation in development that might be attributed to the effect of bleach (Porta-de-la-Riva *et al*., 2012). Following bleach synchronization, approximately 30 embryos were transferred into the wells of 96-well microplates in 50 μL of K medium (Boyd *et al*., 2012). Embryos were exposed to toxicants using four biological replicates and three technical replicates (bleach groups) for a total of 12 wells per strain in each toxicant condition and experimental block. Embryos were exposed to the two control conditions, DMSO and water, using eight biological replicates and three technical replicates for a total of 24 wells per strain in each control condition and experimental block. The positions of the replicate wells were randomized across the microplates to control for any well position effects. After the embryos were added, microplates were labeled and sealed with gas-permeable sealing film (Fisher Cat #14-222-043), placed in humidity chambers, and incubated overnight at 20°C with shaking at 170 rpm. The following morning, food was prepared for the developmentally arrested first stage larval animals (L1s) using frozen aliquots of HB101 *E. coli* (**Section 2.3**). HB101 aliquots were thawed at room temperature, combined into a single vessel, diluted to OD600 30 with K medium, and treated with kanamycin (final concentration 50 μM) to inhibit further bacterial growth and prevent contamination. Experimental treatments were prepared by adding toxicants or vehicle controls to the food suspension. For each block, all experimental treatments were prepared sequentially in random order and used immediately after preparation. All toxicant conditions were prepared at 3X concentration by thawing a frozen aliquot of toxicant stock solution, diluting it to a working concentration using DMSO or water if necessary, then mixing the required volume of the toxicant solution with OD600 30 HB101 food suspension in a sterile vessel. Depending on the toxicant, DMSO or water was added to ensure the final diluent concentration was 1% of the total volume. The synchronized L1 nematodes were simultaneously fed and treated by adding 25 μL of the 3X concentrated treatment suspension to the 50 μL of K medium in the microplate wells using a 12-channel micropipette. Immediately after feeding, the microplates were sealed with a gas-permeable sealing film (Fisher Cat #14-222-043), returned to the humidity chambers, and incubated for 48 hours at 20°C with shaking at 170 rpm. Afterwards, the microplates were removed from the incubator and all wells were treated with 40.6 mM sodium azide (325 μL of 50 mM sodium azide in 1X M9) to paralyze and straighten the animals. After 10 minutes in sodium azide, well images were acquired using a Molecular Devices ImageXpress Nano microscope (Molecular Devices, San Jose, CA) with a 2X objective. The images were then used to measure toxicant response phenotypes.

### 2.6 Image processing, data cleaning, and toxicant susceptibility calculations

To rapidly extract toxicant response phenotypes from our well images, we wrote a Nextflow (v20.01.0) analysis pipeline that runs parallel instances of CellProfiler (v4.2.0) on the Quest High-Performance Computing Cluster (Northwestern University) (Di Tommaso *et al*., 2017; Stirling *et al*., 2021). The cellprofiler-nf (v1.1.0) workflow can be found at https://github.com/AndersenLab/cellprofiler-nf/tree/v1.1.0. At the core of this pipeline is CellProfiler’s Worm Toolbox, which contains modules that facilitate the extraction of morphological features of individual *C. elegans* animals from images (Wählby *et al*., 2012). In brief, our pipeline identifies regions of interest (ROIs) in an image by segmenting them from the background via intensity thresholding. CellProfiler then uses the UntangleWorms module to find “nematode objects” within the ROIs using four probabilistic nematode models that we trained on similar data (Nyaanga *et al*., 2021; Wählby *et al*., 2012). The morphological features of the segmented ROIs and nematode objects found within the ROIs are then extracted using downstream measurement modules. The final output of our pipeline includes (1) a single R data file with image metadata, assay metadata, nematode object morphology measurements, and the parent ROI measurements for every nematode object found in the image set, and (2) a directory of diagnostic images that show the outlines of all the ROIs and nematode objects found in each image overlaid on the raw image. We ran the cellprofiler-nf pipeline on our experimental images and then cleaned and summarized the data using the *easyXpress* R package (v1.0.0) (Nyaanga *et al*., 2021). The data processing steps are summarized in the paragraph below, and a more detailed description is included in the supplemental material (**Methods S1**).

To clean potentially spurious measures from the raw data, we removed small objects (length < 100 µm), objects near the well edge, and objects for which multiple nematodes were detected within an ROI. For conditions that contained few or no small nematodes, we removed putatively spurious small objects < 165 µm. We then excluded all remaining objects that were classified as “non-nematode” by a stochastic gradient boosted classifier that we trained to 89.6% accuracy with over 1,000 nematode objects from a broad range of experimental conditions using the *caret* R package (v6.0-92) (Kuhn, 2008). Finally, within each well, we removed nematode objects with outlier lengths using Tukey’s fences (Tukey, 1977). To summarize the cleaned data for each well, we calculated the total number of nematodes and the median animal length within each well. We then filtered the summarized well data by removing wells with fewer than five or more than 30 animal length measurements or wells with outlier median animal length values for any combination of strain and condition. Next, we normalized our data by subtracting the average median animal length in control conditions for each strain in each bleach from the median animal length in treatment wells for the same strain. This normalization allowed us to remove the effect of strain differences that we observed in control conditions, estimate strain-specific toxicant susceptibilities, and retain the effects of bleach in our data. Next, within each treatment condition, we estimated the effect size of each bleach using a linear model and removed bleaches with extreme effects. We then regressed out the remaining bleach effects by taking the residual values from a linear model of bleach effects that excluded the bleaches with extreme effects. We refer to these final, zero-centered residual values as “normalized length differences” and use them to analyze toxicant susceptibilities throughout the manuscript.

Finally, we filtered all data for a strain in a particular treatment if fewer than three wells were retained after applying the filters above.

### 2.7 Heritability calculations

The total phenotypic variance (*VP*) of a given toxicant response can be partitioned into genetic variance (*VG*) and environmental variance (VE) components (*VP = VG*+VE). Broad-sense heritability (*H*^2^) describes the proportion of phenotypic variation attributable to genetic variance. To estimate *H*^2^ for each toxicant response, we fit a linear mixed-effects model to the phenotype data with strain coded as a random effect using the *lme4::lmer* function from the *lme4* (v1.1.27.1) package in R (Bates *et al*., 2015). We then extracted the effect of strain (*VG*) and the residual variance (*VE*) from the model and calculated *H*^2^ with the equation *H^2^ = VG / (VG*+VE). We estimated confidence intervals around the *H*^2^ estimates using a parametric bootstrap approach implemented with the *lme4::bootMer* function from the *lme4* (v1.1.27.1) package in R. We also estimated narrow-sense heritability (*h^2^*) or the proportion of phenotypic variation (*VP*) attributable to just additive genetic variance (*VA*), using a previously described method (Widmayer, Crombie, *et al*., 2022). Briefly, we made a genotype matrix for the GWAS strains using the hard-filtered variant call format (VCF) file from CaeNDR (Release 20220216) (Crombie *et al*., 2024). We then calculated the variance-covariance matrix (*MA*) from the genotype matrix using the *sommer::A.mat* function from the *sommer* (v4.1.5) R package (Covarrubias-Pazaran, 2016). Next, we estimated *VA* using the linear mixed-effects model function *sommer::mmer* with strain as a random effect and *MA* as the covariance matrix. Finally, we calculated *h^2^* and its standard error using the *sommer::vpredict* function.

### 2.8 Correlations between toxicant susceptibilities and the identification of extreme responders

To assess the consistency of strain susceptibilities to multiple toxicants, we conducted a pairwise correlation analysis. We used the *psych::corr.test* function from the *psych* (v2.4.12) package in R to calculate Spearman’s rank correlation coefficients and applied a Bonferroni correction to adjust *p*-values. We visualized the correlation results and performed hierarchical clustering of the correlation matrix with the *pheatmap::pheatmap* function from the *pheatmap* (v1.0.12) package from R. We examined whether differences in susceptibility were more correlated when condition pairs involved toxicants from the same group (use or MoA) by estimating the average correlation of within-group condition pairs and between-group condition pairs. To identify individuals with susceptibility that significantly deviated from the population mean, we applied a threshold of two standard deviations (SD). For each toxicant condition, we calculated the mean and SD of normalized length differences of all individuals. We then classified individuals with responses exceeding the mean normalized length difference by more than two SDs as minimally susceptible, and those with responses below the mean by more than two SDs as highly susceptible.

### 2.9 Genome-wide association (GWAS) mappings

We applied the NemaScan (commit hash: b58711369124885fb90ce9c53b720313fa68f79b, container id: andersenlab/nemascan:20220407173056db3227) GWAS mapping pipeline (Widmayer, Evans, *et al*., 2022) to identify susceptibility-associated genomic regions for all toxicant response traits. We conducted association mapping with biallelic SNVs from the *C. elegans* hard-filtered variant call format file (VCF) on CaeNDR (Release 20220216) (Crombie *et al*., 2024). The VCF was filtered to include SNVs with no missing genotypes and a minor allele frequency (MAF) of 5% in the mapping panel. To reduce redundancy and control for markers in linkage disequilibrium (LD), we further pruned SNVs using PLINK v1.9 (Chang *et al*., 2015; Purcell *et al*., 2007) with the *–indep-pairwise 50 10 0.8* option. The remaining 14,852 SNVs served as markers in the association mapping. To test markers for association with individual differences in susceptibility, we employed two mapping algorithms from the Genome-wide Complex Trait Analysis (GCTA) software package (Yang *et al*., 2011): *–fastGWA-lmm-exact* (Inbred) and *–mlma-loco* (LOCO). Each algorithm used a sparse genetic relatedness matrix (GRM) to account for the relatedness among strains within a mixed-effects linear model (Yang *et al*., 2011). We constructed sparse matrices for the Inbred and LOCO (Leave-One-Chromosome-Out) mapping algorithms with *–make-grm-Inbred* (designed for inbred organisms) and the *–make-grm* method, respectively. The Inbred algorithm adjusts the genetic relatedness matrix to more accurately model inbred populations (Jiang *et al*., 2019), whereas the LOCO algorithm excludes variants on the chromosome containing the focal SNV from the relatedness matrix to prevent confounding by local linkage (Yang *et al*., 2014). To further control the potential increase in false discoveries caused by residual population structure, we performed eigendecomposition on the GRMs and included the first eigenvector as a covariate in the linear mixed model.

We performed eigendecomposition of the genotype matrix of marker SNVs to estimate the number of independent markers and used this value to define the eigen-based significance threshold. For each mapping approach, we defined susceptibility-associated regions by aggregating neighboring significant markers into a single region of interest. To account for extensive LD, each region was extended by an additional 50 markers. Finally, we merged regions associated with differences in susceptibility that were separated by fewer than 200 markers into a single region of interest. We identify each susceptibility-associated region by the chromosome and position of the most significant SNV (*e.g*., V:1486141), referred to as the peak marker. We evaluated concordance between the GWAS methods (Inbred and LOCO) by comparing the susceptibility-associated regions that each algorithm detected. We detected a total of 79 regions: 40 with Inbred and 39 with LOCO. 42 susceptibility-associated regions (Inbred = 22, LOCO = 20) were detected exclusively by one method. The other regions were identified by both methods, meaning the associated genomic regions physically overlapped. Simulations of *C. elegans* GWAS show no significant difference in detection power or false discovery rates between the two methods (Widmayer, Evans, *et al*., 2022). However, mappings performed with LOCO have higher genomic inflation, suggesting that the Inbred algorithm provides a better correction for population structure than LOCO (Widmayer, Evans, *et al*., 2022). Therefore, we focused on the susceptibility-associated regions detected using the Inbred approach.

### 2.10 Estimating the effect sizes and the proportion of variation explained by susceptibility-associated regions

We estimated two parameters to characterize each susceptibility-associated region: the phenotypic effect size and the proportion of phenotypic variance explained. To estimate phenotypic effect size, we extracted the *β* coefficient from the linear mixed model used in the GWAS, which represents the change in normalized length differences between strains with differing alleles at the peak marker. To calculate the proportion of phenotypic variance explained by each region, we first fit a one-way ANOVA with normalized length difference as the response variable and allelic state at the peak marker for the region as the predictor. The proportion of phenotypic variance explained by each region was calculated as the ratio of the sum of squares for peak marker genotype (*SSallele*) to the total sum of squares (*SStotal*) from the ANOVA model (*var.exp = SSallele / SStotal*).

### 2.11 Identifying biological processes over-represented in susceptibility-associated regions

For each toxicant, we conducted a Gene Ontology (GO) term over-representation analysis on genes within each susceptibility-associated region (Boyle *et al*., 2004). First, we filtered the WormBase gene feature file (GFF) (PRJNA13758, WS283) (Sternberg *et al*., 2024) to include only genes entirely contained within each susceptibility-associated region (*i.e.*, genes where both the start and stop coordinates fell within interval boundaries). We then applied the *clusterProfiler::enrichGO* function from the *clusterProfiler* (v4.6.2) package in R (Wu *et al*., 2021) to test for over-representation of biological processes among genes contained within susceptibility-associated regions. To control for multiple hypothesis testing, we set a Bonferroni-adjusted *p-*value significance threshold of 0.05 and a *q*-value threshold of 0.05.

### 2.12 Identification of candidate genes from susceptibility-associated genomic regions

To identify the genes that are most likely to contribute to individual differences in susceptibility, we performed a fine-mapping analysis on each susceptibility-associated region identified by the GWAS. Fine mapping tests all biallelic SNVs within a region for association with susceptibility using the Inbred algorithm, rather than the limited set of marker SNVs used for GWAS. Candidate genes were identified in the fine-mapping data by focusing on genes containing SNVs that had *p*-values above a significance threshold (-log10(*p*-value) > 5), were not rare (> 0.05 MAF), were in moderate to strong LD with the peak GWAS marker (*r*^2^ > 0.75), and were predicted to have high impact on gene function. We used the Variant Annotation tool from CaeNDR (Crombie *et al*., 2024) to classify the predicted consequences of each SNV on gene function as either high-impact (affects amino acids, start/stop codons, or splice sites) or as low-impact (all other variants). LD was measured between each SNV from the fine mapping and the peak marker from the GWAS using PLINK v1.9 (Purcell *et al*., 2007). To summarize the biological processes associated with the candidate genes identified through fine mapping, we assigned GO terms to each candidate gene with the *clusterProfile::groupGO* function (level = 5) from the *clusterProfiler* R package (v4.6.2) (Wu *et al*., 2021). We also performed a GO term over-representation analysis on the candidate gene set using the same method previously applied to the full gene sets from susceptibility-associated regions (**Section 2.11**).

### 2.13 Physical overlaps and LD between susceptibility-associated regions

We identified physical overlaps between susceptibility-associated regions for each pair of toxicant response traits. For each trait pair, we quantified the proportion of overlapping regions relative to the total number of regions identified for both traits using the formula: *(Regionsoverlapping/(RegionsA+RegionsB)*. To examine whether any two overlapping regions were in LD, we calculated the pairwise correlation coefficient (*r*^2^) between peak markers from each region using PLINK (v1.9) (Purcell *et al*., 2007).

### 2.14 Semantic similarity of genes associated with differences in susceptibility

For each toxicant condition pair, we measured the functional similarity of genes within susceptibility-associated regions with GO term semantic similarity analysis. We used the *GoSemSim::mClusterSim* function from the *GoSemSim* (v2.24.0) package in R (Yu *et al*., 2010) to calculate the functional similarity of genes from the graph structure of the GO terms (Wang *et al*., 2007). We conducted this analysis with two gene sets: all genes within any susceptibility associated regions for each toxicant condition, and with candidate genes identified within these regions. We restricted our analysis to gene features from the WormBase GFF (PRJNA13758, WS283) (Sternberg *et al*., 2024) contained entirely within all susceptibility-associated regions detected for each toxicant. Biological process GO terms were sourced from the org.Ce.eg.db database (v3.16.0) (Carlson, 2019) after converting gene IDs from Ensembl to Entrez with the *clusterProfiler::bitr* function from the *clusterProfiler* R package (v4.6.2) (Wu *et al*., 2021). Semantic similarity analysis was performed with the *GoSemSim::mClusterSim* function using the best-match average (BMA) method, which specifies how to handle genes with multiple GO terms. To visualize results and perform hierarchical clustering of toxicant traits by gene content similarity, we used the *pheatmap::pheatmap* function from the *pheatmap* (v1.0.12) package in R.

### 2.15 Prioritization of susceptibility-associated regions for additional follow-up

We manually reviewed all susceptibility-associated regions and developed a prioritization scheme by considering the following criteria. First, we excluded regions where fewer than 20 strains carried the minor allele at the peak marker. This criterion ensures that a sufficiently large set of phenotypically and genetically distinct individuals were available to validate effects and narrow regions of interest using subsequent crosses (Andersen and Rockman, 2022). Next, we implemented a prioritization scheme to rank the regions that remained (n = 22) from most actionable (1) to least actionable (22) by considering the following characteristics. The first metric in our prioritization scheme was the significance of association. Susceptibility-associated regions with higher peak marker -log10(*p*-values) were prioritized above those regions with less significant signatures of association. The second metric in our prioritization scheme was the extent of LD within each susceptibility-associated region. Large genomic regions of interest, with many variants in high LD, are more challenging to narrow to single-gene resolution because of their physical size and the strong linkage between alleles (Andersen and Rockman, 2022). Therefore, we prioritized smaller genomic regions where the signature of association was localized around the peak marker. We classified fine mapping plots of susceptibility-associated regions into one of three categories (Less than 250 kb, 250 to 500 kb, or Greater than 500 kb) by considering the physical distance of significant markers in high LD (*r^2^* > 0.5) with the peak marker and prioritized regions categorized as Less than 250 kb or 250 to 500 kb. The final criterion was the overlap of each susceptibility-associated region with regions of extreme genetic diversity or hyper-divergent regions (HDRs). HDRs are shared by subsets of strains and often differ in gene content relative to the reference genome (Thompson *et al*., 2015; Kim *et al*., 2019; Lee *et al*., 2021; Moya *et al*., 2024). Because the exact genomic content of HDRs is unknown, a high incidence of HDRs within a susceptibility-associated region can complicate candidate gene identification. We located HDRs in each strain using features of short-read sequencing data (**Methods S2**), calculated the proportion of strains with at least one HDR overlapping each susceptibility-associated region, and prioritized regions where fewer strains carried overlapping HDRs. Human orthologs for candidate genes were identified using data from ortholist2 (Kim *et al*., 2018).

## 3 Results

### 3.1 Toxicant susceptibilities differ widely across 195 *C. elegans* wild strains

We used a high-throughput phenotyping platform to quantify the differences in susceptibility among 195 wild *C. elegans* strains. Synchronized first-larval-stage animals were exposed to each toxicant for 48 hours in 96-well plates (**Section 2.5, Table S2**). In control conditions, animals increase in length continuously as they develop (Knight et al. 2002; Nyaanga et al. 2022). However, exposure to toxicants reduces growth, and the magnitude of inhibition varies among strains. Growth inhibition was quantified for each strain as the average difference in mean body length between toxicant and control wells, thereby normalizing for inherent growth differences among strains. We then used zero-centered normalized length differences, which are residuals from a regression correcting for bleach effects (**Section 2.6**), to compare susceptibility across strains for a given toxicant (**Table S3**). Strains with negative normalized length differences exhibited greater growth inhibition (higher susceptibility), whereas strains with positive values were less affected by the toxicant.

We found large differences in susceptibility among strains to particular toxicants (**Figure S2, Table S4**). For example, normalized length differences ranged from -198 μm to 162 μm in mercury, reflecting large differences in the growth inhibition experienced by the most susceptible strain (ECA396) and the least susceptible strain (JU1934) (**Figure S2**). In total, 108 strains displayed extreme susceptibilities relative to the rest of the population in at least one toxicant condition. High susceptibility (normalized length differences greater than two standard deviations below the mean) to a toxicant condition was more prevalent than minimal susceptibility (normalized length differences greater than two standard deviations above the mean) (**Figure S3**, **Table S4**). The extent of differences in susceptibility among strains varied across toxicant conditions. The standard deviation (SD) of normalized length differences, which represents how much the average strain differed in susceptibility from the mean, ranged from ± 18 μm for silver 7.8 μM, the least variable condition, to ± 70 μm for mercury, the most variable condition (**Figure S2**, **Table S4**). Notably, the variability in normalized length differences was generally similar across toxicant use groups (**Table S5**). Overall, the variability in toxicant susceptibilities suggests that genetic differences among the strains contribute to these responses.

### 3.2 Genetic differences among wild strains influence toxicant susceptibilities

To evaluate how genetic differences influence toxicant susceptibilities, we estimated broad-sense heritability (*H*^2^) and narrow-sense heritability (*h*^2^) for each toxicant condition (**Figure 1)**. Broad-sense heritability (*H*^2^) reflects the proportion of variation in susceptibility explained by genetic differences among strains, including both individual variants with additive effects and non-additive genetic interactions between variants (such as epistatic and dominance effects). On average, these genetic factors collectively explained more than half the variance in susceptibility among strains (mean toxicant condition *H*^2^ = 0.53 ± 0.16 SD; **Figure 1, Table S6**). Narrow-sense heritability (*h*^2^), which captures only the proportion of variance in susceptibility attributable to additive genetic effects alone, was expectedly lower for all toxicant conditions (mean *h*^2^ = 0.21 ± 0.13 SD; **Figure 1, Table S6**). Importantly, the *h*^2^ estimates are representative of what can be detected using a GWAS, which assumes all variants contribute to differences in susceptibility additively and does not model non-additive interactions (Yang *et al*., 2011). Ultimately, the high heritability estimates described here (mean *h*^2^ > 0.2) suggest that GWAS will have sufficient statistical power to detect genomic regions associated with differences in toxicant susceptibility among strains.

**Figure 1.**
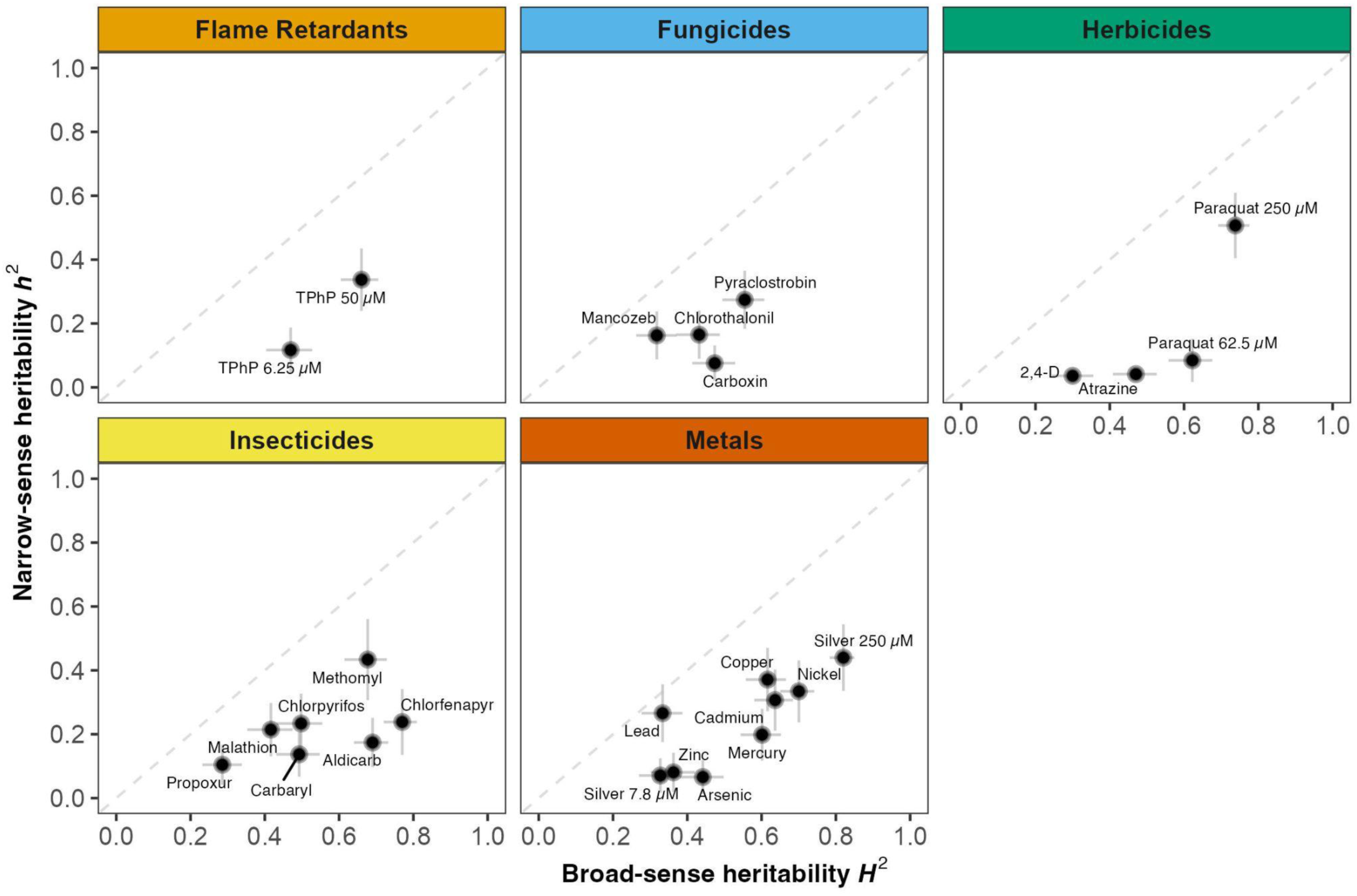
Heritability of toxicant susceptibilities. The broad-sense (*H^2^*) (x-axis) and narrow-sense heritability (*h^2^*) (y-axis) of normalized length differences were calculated for each toxicant exposure condition. Within each use group, heritability estimates for a single toxicant exposure condition are denoted by a cross, where the horizontal line corresponds to the 95% confidence interval of the broad-sense heritability estimate obtained using the parametric bootstrapping approach, and the vertical line corresponds to the standard error of the narrow-sense heritability estimate. The three toxicants measured at two concentrations (TPhP, paraquat, and silver) are annotated with concentrations.

The large proportion of differences in susceptibility attributable to genetic factors likely reflects our selection strategy for toxicant conditions. We specifically chose toxicant concentrations where differences in susceptibility had the highest heritability across a small panel of eight genetically diverse strains (Widmayer, Crombie, *et al*., 2022). From this prior assay, where multiple concentrations were tested for each toxicant, we selected the most heritable concentration for each toxicant to maximize our ability to detect specific genetic variants using GWAS. We assumed that genetic variation observed in the small panel of genetically diverse strains would reflect the heritability of toxicant susceptibilities across a larger population. To evaluate this assumption, we compared the heritability estimates made here using 195 strains with those estimates derived previously using just eight strains. We found significant positive correlations between the previous heritability estimates and our current heritability estimates across all 26 toxicant conditions (**Figure S4**). These correlations demonstrate that preliminary screens of toxicity across a small panel of diverse strains can provide insights into the heritability of differences in susceptibility across a larger population.

### 3.3 Strain response correlations point to mechanisms influencing susceptibility

Organisms rely on cellular stress response pathways to mitigate the harmful effects of toxicant exposures. In *C. elegans*, some of these pathways act generally across multiple toxicants (Blackwell *et al*., 2015; Labbadia and Morimoto, 2015; Shore and Ruvkun, 2013). By contrast, others are more specific. For example, the xenobiotic detoxification pathway contains proteins that target specific compounds, such as toxicants with similar chemical properties or mechanisms of action (MoA) (Hartman *et al*., 2021). We hypothesized that shared genetic variation could affect general aspects of stress response pathways and lead to similar susceptibility patterns across toxicant conditions. To test this hypothesis, we assessed how similarly strains responded to different toxicants by calculating pairwise Spearman’s rank correlation coefficients of normalized length differences (**Section 2.9**) (Spearman, 1904). These coefficients (ρ) reflect the degree to which strain susceptibility rankings are consistent across toxicant pairs. Most condition pairs showed weak to moderate positive correlations (mean ρ = 0.189 ± 0.152 SD), indicating that on average strains respond somewhat similarly across exposures (**Figure 2**). Of the 325 possible toxicant condition pairs, 88 showed significant correlations, all of which were positive (Bonferroni-corrected *p*-value ≤ 0.05) (**Figure 2**). These correlations were not driven by a few extreme strains, because strains rarely displayed extreme responses across multiple toxicants (**Figure S3, Table S4**). To examine broader response patterns, we performed hierarchical clustering on the correlation matrix, which grouped toxicants by similarity in strain-level responses (**Section 2.8**). The largest of two main clusters contained 65% (17/26) of the toxicant conditions, suggesting that shared genetic variation in general response pathways might influence susceptibility to multiple toxicants (**Figure 2**). However, the overall correlation structure is weak, suggesting that toxicant-specific mechanisms also contribute to variation in susceptibility.

**Figure 2.**
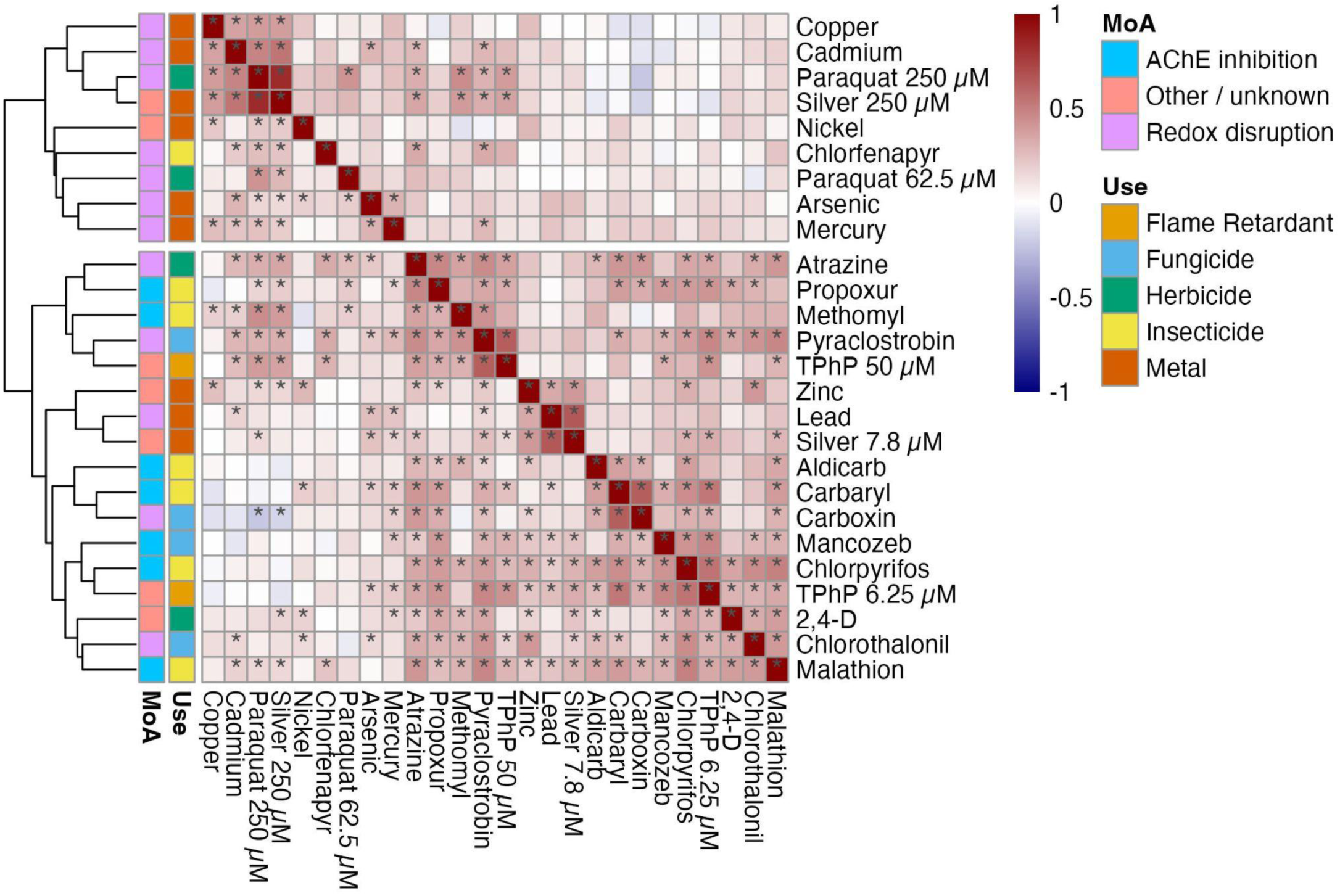
Correlations between differences in susceptibility to toxicant conditions. The pairwise Spearman correlation coefficient (ρ) between all toxicant conditions is displayed as a correlation matrix and colored as a heatmap. Spearman’s rank order correlation coefficients (colored cells), were calculated using the normalized length differences for all strains. Each cell represents the correlation between two toxicant conditions. The main diagonal shows self-correlations (ρ = 1) and divides the upper and lower triangles of the symmetrical correlation matrix. Significant correlations (*p*-value ≤ 0.05) are denoted with an asterisk (*). The upper right triangle shows significance symbols after Bonferroni multiple testing correction, and the lower left triangle shows uncorrected significance symbols. On the left, columns of colored tiles denote the mechanism of action (MoA) and class of each toxicant. The traits are arranged by hierarchical clustering performed with correlation coefficients. More similar toxicant conditions are grouped together. The similarity between toxicants is represented by the branch lengths in the dendrograms plotted to the left of the heatmap.

We further tested whether susceptibility correlations were stronger among toxicants with similar use or MoA classifications. Condition pairs from the same use group or MoA did not exhibit markedly stronger correlations than those correlations between different groups. For example, carbaryl (an insecticide) and carboxin (a fungicide) showed one of the highest correlations, despite differing in both use and MoA (ρ = 0.69, *p*-value < 0.0001); **Figure 2**). The average correlation among within-use group pairs (ρ = 0.235 ± 0.137 SD) was only slightly higher than between-use group pairs (ρ = 0.176 ± 0.153 SD). Similarly, within- and between-MoA group pairs had comparable average correlations (ρ = 0.199 ± 0.144 SD vs. ρ = 0.184 ± 0.156 SD, respectively). Consistent with these findings, hierarchical clustering revealed that toxicants from the same use group or MoA were dispersed throughout the largest cluster, with no clear grouping by either criterion. The exception was a subset of redox-disrupting heavy metals (copper, cadmium, arsenic, and mercury), which exhibited stronger within-group correlations (**Figure 2**). The modest correlation structure and lack of grouping by use or MoA suggest shared genetic variation in general stress response pathways could contribute to variation in susceptibility across multiple toxicants. However, the stronger correlations between redox-disrupting heavy metals might indicate that toxicant-specific mechanisms contribute as well.

### 3.4 GWAS detects genomic regions associated with differences in toxicant susceptibility

To more directly assess the genetic basis of susceptibility differences, we performed a GWAS of all toxicant conditions to pinpoint biological processes and candidate genes underlying strain-specific responses. Our GWAS approach used a representative set of 14,852 marker SNVs distributed across the *C. elegans* genome to test for associations with susceptibility to each toxicant (**Section 2.9**). This analysis identified 236 significant markers across 16 of the 26 toxicant conditions (**Figure S5**). Because nearby markers often reflect the same signal of association, we grouped neighboring significant markers into 40 distinct genomic regions associated with susceptibility (**Figure 3, Table S7**). For some analyses, we used the most significantly associated variant within each region as a representative marker, referred to as the “peak marker”. Among the 16 toxicant conditions with significant associations, the number of detected regions ranged from one to eight, with a median of 1.5 (**Table S7**). The absolute effect sizes of the peak markers for each region ranged from 5 to 51 µm in normalized length difference, and the frequency of those variants in the GWAS population ranged from 5.1% to 47.9% (**Figure 4**, **Table S7, Section 2.10**). Notably, peak markers with larger effects tended to occur at lower frequencies (**Figure 4**), consistent with evolutionary expectations that alleles of large effect are maintained at low frequencies by selection (Orr, 2005). For the ten toxicant conditions without significant associations, susceptibility is likely influenced by variants that either cause small effects, are rare, or participate in epistatic interactions, which are more difficult to detect with GWAS (Manolio *et al*., 2009).

**Figure 3.**
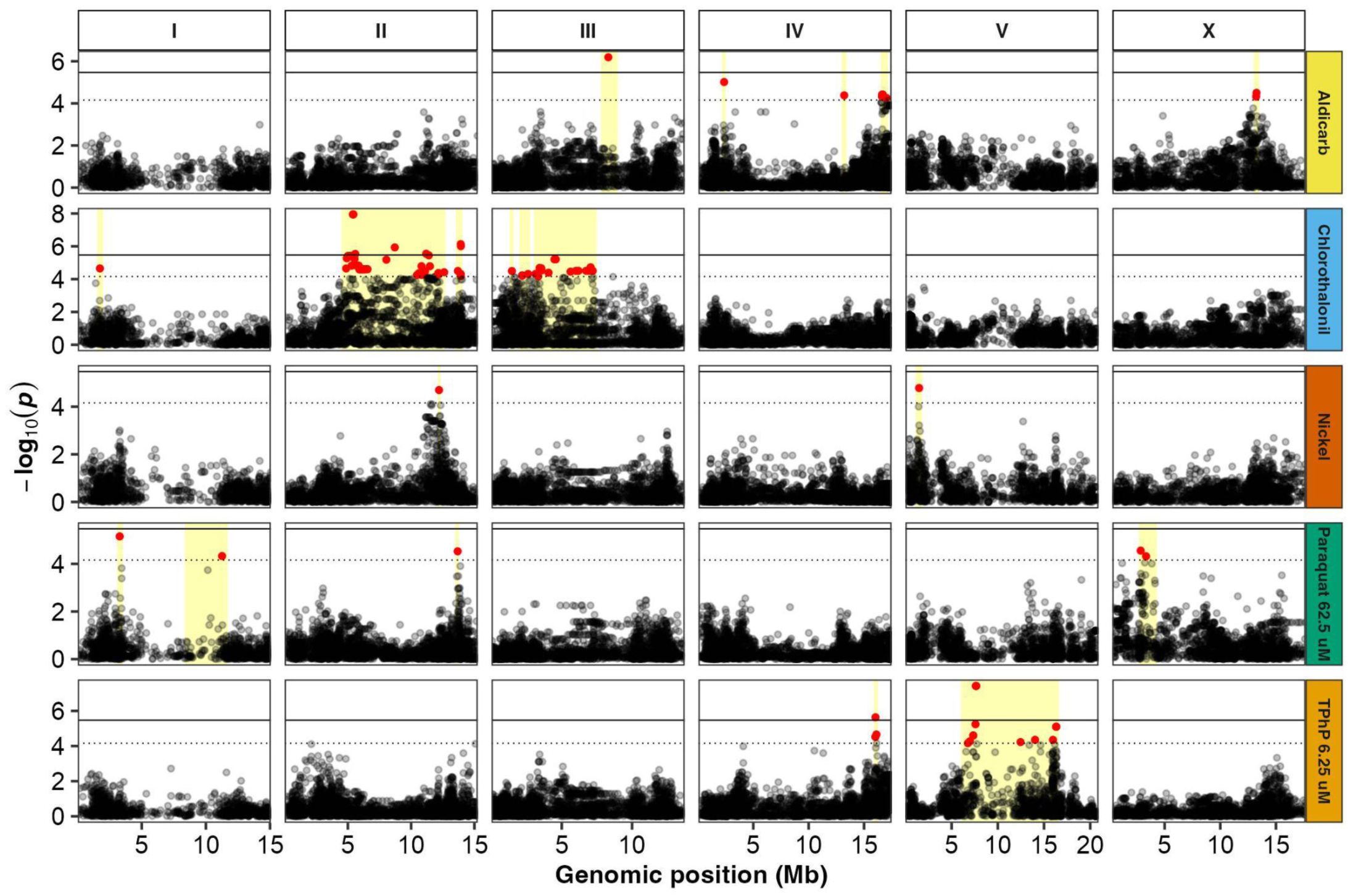
Representative Manhattan plots showing 19 of 40 susceptibility-associated regions identified with GWAS . Representative genome-wide association mappings of normalized length differences for each toxicant use-group are shown. For each toxicant use-group, we selected a representative toxicant condition based on the heritability and the presence of actionable susceptibility-associated regions. Each point represents a single-nucleotide variant (SNV) that is present in at least 5% of the wild strains exposed to each toxicant condition. Genomic coordinates are plotted (x-axis) against the log-transformed *p*-value of the test of association for each SNV (y-axis). The solid black line is the Bonferroni-corrected significance threshold (BF) using all markers. The dashed horizontal gray line is the Eigen-corrected *p*-value threshold using independent markers corrected for LD (genome-wide eigendecomposition significance threshold). Red points represent SNVs above the EIGEN significance threshold, whereas black points represent SNVs below that threshold. Yellow boxes represent distinct susceptibility-associated regions.

**Figure 4.**
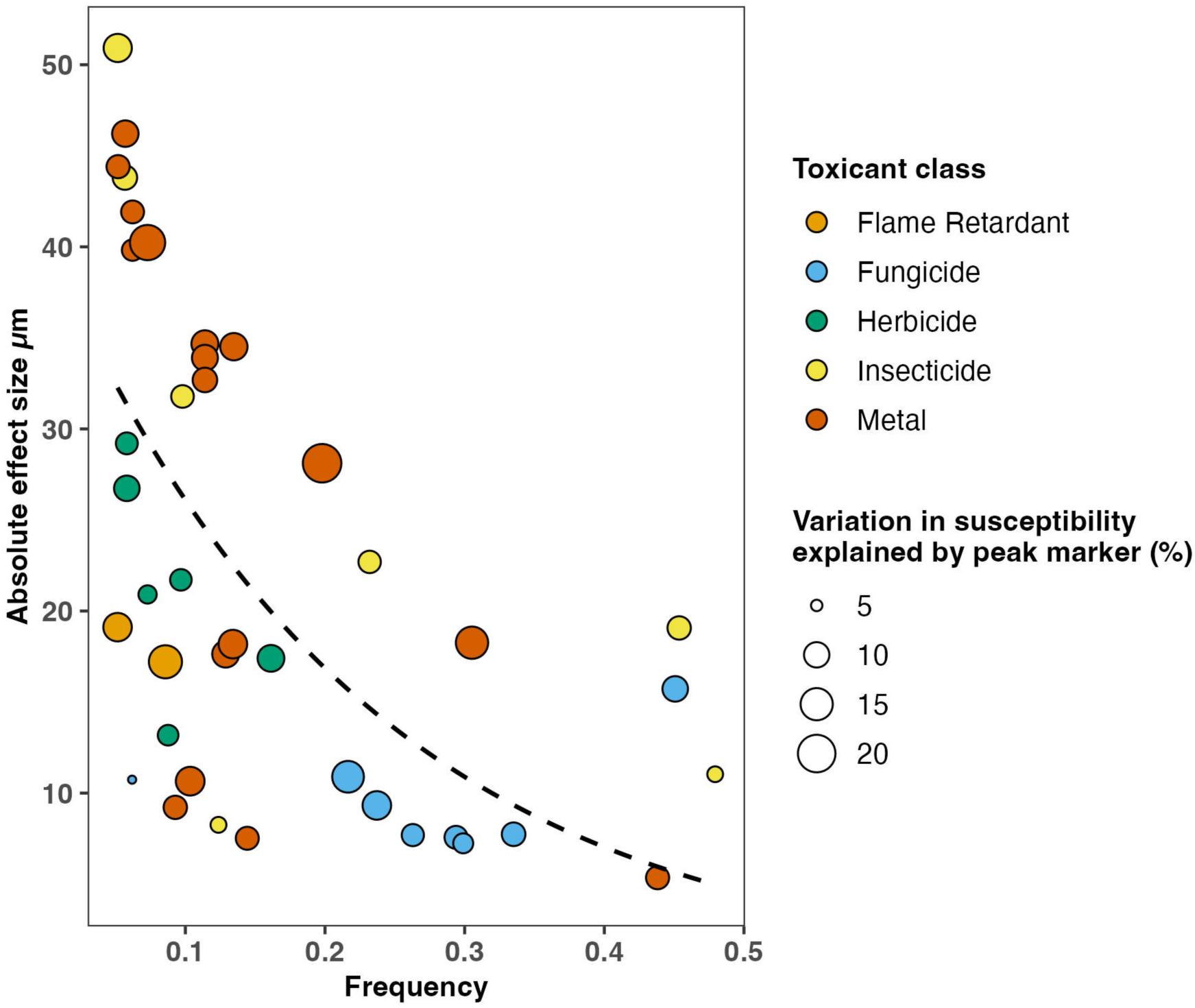
Absolute effect size of susceptibility associated variants by frequency. Each point represents the most significant marker SNV (peak marker) from one of the 40 susceptibility-associated regions, plotted by its frequency in the 195 strains assayed (x-axis) and its absolute effect size (y-axis). The absolute effect size of each peak marker is calculated by taking the absolute value of the effect size estimated by the GWAS linear mixed model (β coefficient, **Section 2.10**). The effect size represents the additive effect of the SNV on the difference in normalized animal length between normalized length differences for strains with different alleles. The point size represents the proportion of variance in normalized animal length differences explained. The point colors indicate the toxicant use group. The dotted line represents the exponential decay fit of the relationship between frequency and absolute effect size for all peak markers.

### 3.5 Functional enrichment and candidate susceptibility genes within GWAS regions

To explore the biological function of 40 susceptibility-associated genomic regions, we analyzed their gene content. Gene ontology (GO) analysis revealed significant enrichment of specific biological processes within 19 susceptibility-associated regions across 11 different toxicant conditions (**Section 2.11**, **Figure S6**). Some of these regions were enriched for genes involved in biological processes known to participate in toxicant responses. For example, three overlapping genomic regions associated with different toxicants (arsenic, II:13692527; chlorothalonil, II:13921653; and paraquat 62.5 µM, II:13675568) were enriched for genes involved in glutathione and sulfur compound metabolism (GO:0006749 and GO:0006790, respectively) (**Figure S6**). Accordingly, nine glutathione-s-transferase (GST) genes, which encode key phase II detoxification enzymes in *C. elegans* (Hartman *et al*., 2021), are in these regions, suggesting that natural variation in detoxification pathways might contribute to differences in susceptibility to these toxicants. A different region associated with mercury (III:3716201) was enriched for ubiquinone biosynthesis genes (GO:0006743, GO:0006744) (**Figure S6**), consistent with reports that mercury disrupts mitochondrial electron transport in *C. elegans* (Ke *et al*., 2023). This region also contains *clk-1*, a conserved gene implicated in mitochondrial stress responses in both *C. elegans* and humans (Monaghan *et al*., 2015; Rodríguez-Aguilera *et al*., 2005). Yet another region (X:2848181) associated with paraquat (62.5 µM) exposure was enriched for carbohydrate and hexose transport genes (GO:0008643 and GO:0008645, respectively), consistent with prior work that demonstrated glucose supplementation alleviates paraquat-induced developmental delays in *C. elegans* and mammalian cell lines (Wang *et al*., 2019) (**Figure S6**). Importantly, other enriched biological processes revealed potential novel connections to toxicant responses.

Not all genes within associated regions are equally likely to influence susceptibility. Genes with variants predicted to impact protein function or transcription are more likely candidates to cause differences in susceptibility than those candidates without functional variation. Using statistical tools and functional annotations for variants, we identified 136 candidate genes across 13 of the 16 toxicant conditions with at least one associated region (**Section 2.12, Table S8**). Several of these candidate genes have plausible connections to toxicant responses. For example, two nuclear hormone receptor (NHR) family members, *nhr-52* and *nhr-121*, were candidate genes for mercury (**Table S8**). NHRs are known regulators of xenobiotic transcriptional responses in *C. elegans (Hartman et al., 2021)*, and elevated expression of *nhr-121* and *nhr-176* has been linked to anthelmintic drug exposure (Dube *et al*., 2023). Additionally, a candidate gene for zinc, *zig-7*, was previously linked to aldicarb resistance in *C. elegans*, which might suggest a potential role in the mediation of multiple toxicant responses (Sieburth *et al*., 2005) (**Table S8**). However, no gene, including *zig-7,* was a candidate for more than one toxicant. This observation contrasts with the modest correlations in susceptibility across toxicants and suggests that variants in distinct genes could underlie responses to different toxicant conditions. Overall, these candidate genes represent compelling targets for future experimental validation.

### 3.6 Differences in susceptibility are associated with distinct genes and biological processes

To further investigate whether the same genes might contribute to differences in susceptibility to multiple toxicants, we examined the physical overlap between susceptibility-associated genomic regions. We found that 70% (28/40) of regions physically overlapped with regions from other toxicant conditions (**Figure 5, Figure S7**).

**Figure 5.**
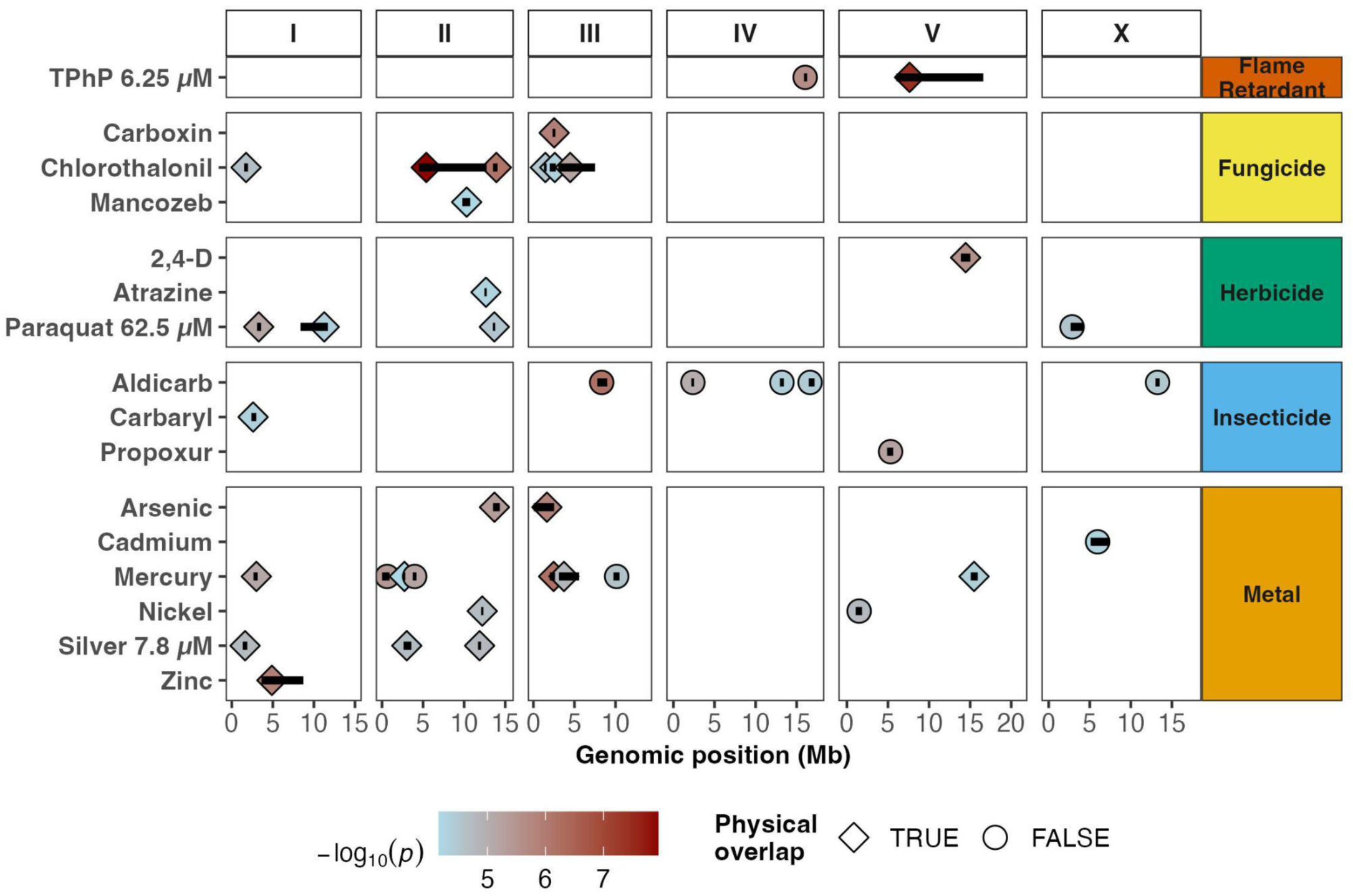
Physical overlaps of genomic regions associated with susceptibility across toxicants. The susceptibility-associated regions for toxicant conditions with at least one marker above the eigen significance threshold are displayed by genomic position (Mb). Each region is represented by a horizontal black bar along its genomic span. The peak marker from each region is shown as a symbol colored according to its significance (-log₁₀(*p*-value)). Diamond-shaped symbols indicate regions that physically overlap with at least one other genomic region associated with a different toxicant condition, whereas circles represent non-overlapping regions. Toxicant conditions are grouped on the y-axis by use classification (Flame Retardant, Fungicide, Herbicide, Insecticide, and Metal) and displayed across the six *C. elegans* chromosomes (I, II, III, IV, V, and X) on the x-axis.

However, these overlaps were limited in scope because most regions overlapped with only one other region (median overlap = 1), and only 16.7% (20/120) of all possible toxicant pairs shared any overlapping regions (**Figure 5**, **Table S9**). Furthermore, we found that the most significant markers from any two overlapping regions showed minimal LD (mean r^2^ = 0.0098, **Section 2.13, Table S9**), supporting the notion that distinct genetic variants within the overlapping regions were associated with differences in susceptibility to each toxicant. Together, these findings demonstrate that, despite some overlaps of susceptibility-associated regions, the same genes are unlikely to influence susceptibility to multiple toxicants.

Although distinct genes were associated with differences in toxicant susceptibility, variants affecting different genes could still contribute to susceptibility differences across multiple toxicants if those genes participate in the same biological pathways. To investigate, we quantified the functional similarity of genes associated with different toxicants by comparing their GO terms (**Section 2.14**, **Figure S8A**) (Wang *et al*., 2007). For each toxicant condition pair, a semantic similarity estimate provides a quantitative measure (ranging from 0 to 1) of how functionally related their associated gene sets are. A high semantic similarity score (near 1) indicates that genes associated with both toxicants are involved in similar biological processes, suggesting the potential of shared mechanisms driving toxicant susceptibility. Conversely, a lower score (near 0) indicates functionally distinct gene sets. Across toxicant condition pairs, the average semantic similarity was 0.675 ± 0.142 (**Figure S87A**), suggesting some functional overlap among gene sets located within susceptibility-associated regions for different toxicants. However, when we refined our analysis to include only candidate genes from those regions, the average functional similarity was much lower (0.245 ± 0.102 SD, **Figure S8B**). Our results show that regions identified by GWAS often contain genes involved in functionally related biological processes, but the specific candidate genes within those regions point to distinct genetic drivers of susceptibility across toxicants.

Identifying the genes that directly influence toxicant susceptibility will require targeted follow-up experiments to validate the effects of each susceptibility-associated region and refine candidate gene lists (Andersen and Rockman, 2022). Some genomic regions are more amenable to approaches used by *C. elegans* researchers for these purposes. Therefore, we prioritized regions that passed our criteria for additional follow-up experiments (**Section 2.15**) and identified 22 regions from 11 different toxicant exposure conditions that included at least one region from every use group, except flame retardants (**Table S10**). Several toxicant conditions had multiple actionable regions, such as chlorothalonil (6), mercury (4), arsenic (2), aldicarb (2), and silver nitrate (7.8 µM) (2) (**Table S10**). The broad representation of toxicants with candidate genes underscores the power of *C. elegans* GWAS for identifying susceptibility-associated regions and candidate susceptibility genes that can be readily validated. Of the 136 candidate genes identified, 94 fell within actionable regions, including 44 with human orthologs (**Table S11**). Natural variation in these genes represents a high-priority target for future experimental validation (**Table S8**).

## 4 Discussion

Our investigation of toxicant responses across 195 genetically diverse *C. elegans* strains revealed substantial natural variation in susceptibility with genetic factors accounting for approximately half of the observed differences among strains (mean *H^2^* = 0.53). Using GWAS, we identified 40 genomic regions associated with differences in susceptibility, encompassing genes in both established toxicant-response pathways and novel biological processes previously unlinked to toxicant susceptibility. We applied prioritization criteria to highlight 22 high-priority genomic regions, which contain 94 candidate susceptibility genes recommended for experimental validation using proven strategies (Andersen and Rockman, 2022). To the best of our knowledge, the majority of the candidate genes (80/94) have not yet been associated with toxicant responses. Moreover, almost half of these genes (44/94) have known human orthologs (Kim *et al*., 2018). These results reinforce the value of *C. elegans* as a robust whole-animal model for the discovery of candidate genes that contribute to differences in toxicant susceptibilities. Ultimately, validation of these candidate susceptibility genes will advance our understanding of the molecular mechanisms that drive toxicant responses with strong translational potential to improve human risk assessments.

Although GWAS is a powerful tool for identifying genetic variants associated with toxicant susceptibility, it cannot capture all the genetic factors influencing these traits. In our study, as in many others, a substantial portion of heritable variation remains unexplained, an observation often referred to as “missing heritability” (Maher, 2008; Mayhew and Meyre, 2017; Rockman, 2012). This gap might come from the inherently low power of GWAS to detect rare variants, common variants with small effects, or non-additive interactions between variants that influence traits like toxicant susceptibility. For example, despite substantial narrow-sense heritability estimates for most toxicant susceptibilities in our study, we did not detect associated variants for 10 of the 26 toxicant conditions. Importantly, the detection of rare variants, common small effect variants, and non-additive interactions among variants is most straightforward in tractable model systems (Bloom *et al*., 2015; Ehrenreich *et al*., 2012), including *C. elegans* using established crossing and mapping schemes (Gaertner *et al*., 2012; Noble *et al*., 2017; Evans *et al*., 2018; Hahnel *et al*., 2018; Evans *et al*., 2021; Andersen and Rockman, 2022; Zhang *et al*., 2022). Future investigations that employ these techniques might identify additional candidate susceptibility genes for the toxicants tested here and other high-priority environmental toxicants. Furthermore, as genomic resources continue to expand, including long-read assemblies that can accurately resolve variation in HDRs (Thomas *et al*., 2015; Lee *et al*., 2021; Stevens *et al*., 2022), future studies can build on this foundation to better characterize the genetic factors that influence toxicant susceptibility and extend these insights across species.

Our prioritization scheme for candidate susceptibility genes is constrained by the resolution of SNV mapping and the assumptions built into our selection criteria. Each region that we identified might harbor insertions, deletions, copy-number, or non-coding variants that influence gene function (Evans and Andersen, 2020). These variants could also drive toxicant susceptibility traits but are undetectable by our current approach, which only considers SNVs and prioritizes SNVs that alter protein-coding sequences. Moreover, LD among nearby variants can obscure causal relationships, as correlated SNVs often share association signals and can mask the presence of multiple causal variants within a region (Cutter and Payseur, 2013). Ultimately, integrating complementary data, including gene expression profiles and structural variant calls, together with mediation analyses (MacKinnon *et al*., 2007) that model causal relationships among genotype, gene expression, and phenotype (Evans and Andersen, 2020; Zhang *et al*., 2022), will be critical for capturing the full spectrum of variants contributing to toxicant susceptibility.

Future work can translate *C. elegans* discoveries to human toxicology. First, the candidate susceptibility genes described here need to be functionally validated using experimental crosses and genetic tools such as CRISPR-Cas9 genome editing to confirm their role in toxicant response. Once validated, the human orthologsof these genes can be identified using established orthology databases (Kim *et al*., 2018; Emms and Kelly, 2019). Population-scale human genomic datasets can then be used to characterize natural genetic variation in these orthologs across diverse human populations (Chen *et al*., 2024; Byrska-Bishop *et al*., 2022). Next, variants with predicted functional impacts can be introduced into CRISPR-edited human cell lines to evaluate their impacts on toxicant responses. Ultimately, the discovery of functionally important variants in human models can support the development of susceptibility biomarkers and improve chemical risk assessment by accounting for human genetic diversity.

## 5 Acknowledgements

We would like to thank members of the Andersen laboratory for their feedback and helpful comments on this manuscript. We thank the *C. elegans* Natural Diversity Resource (NSF Capacity grant 2224885) for providing us with strains for this study. This work was supported by an NIH NIEHS grant (ES029930) to E.C.A. and M.V.R.R.M. was supported by the National Institutes of Health Training Grant (T32 GM008449) through Northwestern University’s Biotechnology Training Program.

## 6 Data availability

The data and code required to perform the analyses in this paper are maintained in a GitHub repository (https://github.com/AndersenLab/ToxinGWAS_Manuscript/tree/master).

## AUTHOR CONTRIBUTIONS

**Conceptualization:** E.C.A

**Data curation:** R.M, T.A.C, N.D.M, A.O.S, J.B.C

**Formal analysis:** R.M, T.A.C

**Funding acquisition:** E.C.A

**Investigation:** R.M, T.A.C, S.J.W, J.W, J.B.C, R.E.T

**Methodology:** R.M, T.A.C, S.J.W, J.W, J.B.C

**Project administration:** E.C.A

**Resources:** E.C.A, R.E.T, C.B., L.S, L.V.S, M.V.R, M.G.S, M.-A.F

**Software:** T.A.C, S.J.W, R.M

**Supervision:** E.C.A

**Validation:** T.A.C, R.M

**Visualization:** T.A.C, S.J.W, R.M

**Writing - original draft:** T.A.C, R.M

**Writing - reviewing & editing:** T.A.C, R.M, S.J.W, E.C.A, A.O.S, J.W.

## Notes

https://github.com/AndersenLab/ToxinGWAS_Manuscript

### Competing Interest Statement

The authors have declared no competing interest.

