## Supplemental Materials for "Natural variation suggests candidate genes underlying *Caenorhabditis elegans* susceptibility to diverse toxicants"

**Figure S1.** GWAS simulations to optimize mapping panel strain selection

**Figure S2.** Differences in susceptibility among 195 *C. elegans* strains to 26 environmental toxicant conditions

**Figure S3.** Number of strains with extreme susceptibility profiles across 26 toxicant exposures

**Figure S4.** Regression of heritability estimates for the 195 strains in the GWAS mapping panel against the eight strains from Widmayer et al. 2022

**Figure S5.** Susceptibility-associated intervals for all 26 toxicant exposures

**Figure S6.** Gene ontology enrichment of susceptibility-associated genomic intervals

**Figure S7.** Physical overlaps of susceptibility-associated intervals across toxicants

**Figure S8.** GO term semantic similarity of susceptibility-associated genes

**Table S1.** Strains selected for the GWAS panel

**Table S2.** Environmental toxicants and exposure conditions assayed

**Table S3.** Strain-specific toxicant response data

**Table S4.** Differences in susceptibility to toxicant conditions among strains

**Table S5.** Comparison of the mean standard deviation of normalized animal lengths across toxicant groups

**Table S6.** Heritability estimates for toxicant conditions

**Table S7.** Susceptibility-associated intervals detected by the Inbred GWAS algorithm across toxicant conditions

**Table S8.** Candidate genes from susceptibility-associated intervals detected by GWAS

**Table S9.** Overlapping susceptibility-associated intervals across toxicant condition pairs

**Table S10.** Prioritization of actionable susceptibility-associated regions detected by GWAS

**Table S11.** Human orthologs for candidate genes

**Methods S1.** Detailed image processing and data cleaning steps

**Methods S2.** Identification of hyper-divergent regions in *C. elegans*

Supplementary Tables

### SUPPLEMENTAL FIGURES

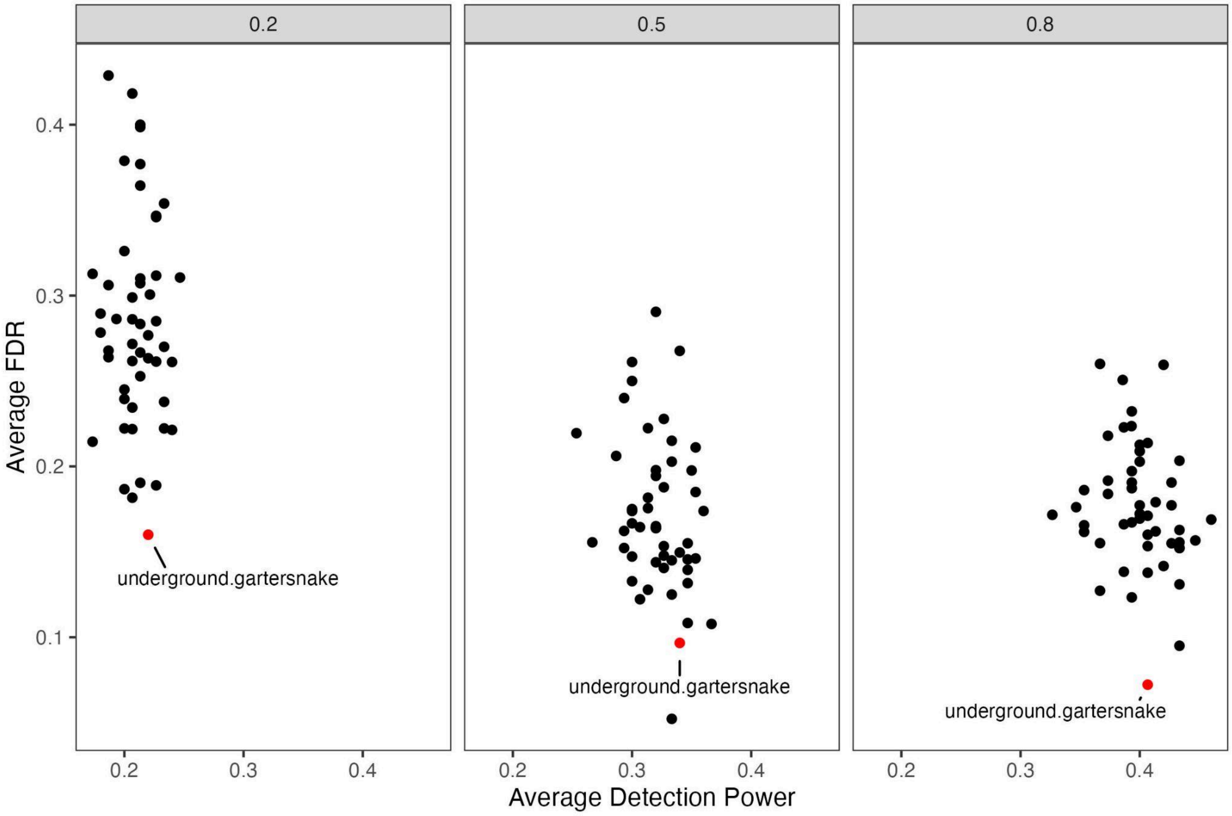


**Figure S1. GWAS simulations to optimize mapping panel strain selection.** The average power (proportion of simulated susceptibility-associated intervals detected) and the average false discovery rates (FDR) (proportion of detected susceptibility-associated intervals that were not simulated) for 30 GWAS simulations with 50 unique 192 strain mapping panels. The mapping panels were generated by randomly sampling 181 genetically related (swept) strains to pair with the 11 strains included in previous dose-response publications. A point represents the average performance of each mapping panel across 30 simulated GWAS. The red colored point represents the simulated performance of the strains included in the GWAS and was assigned the label "underground.gartersnake". The performance of each strain set is faceted by the narrow-sense heritability of the simulated trait ($h^2$ = 0.2, 0.5, 0.8).

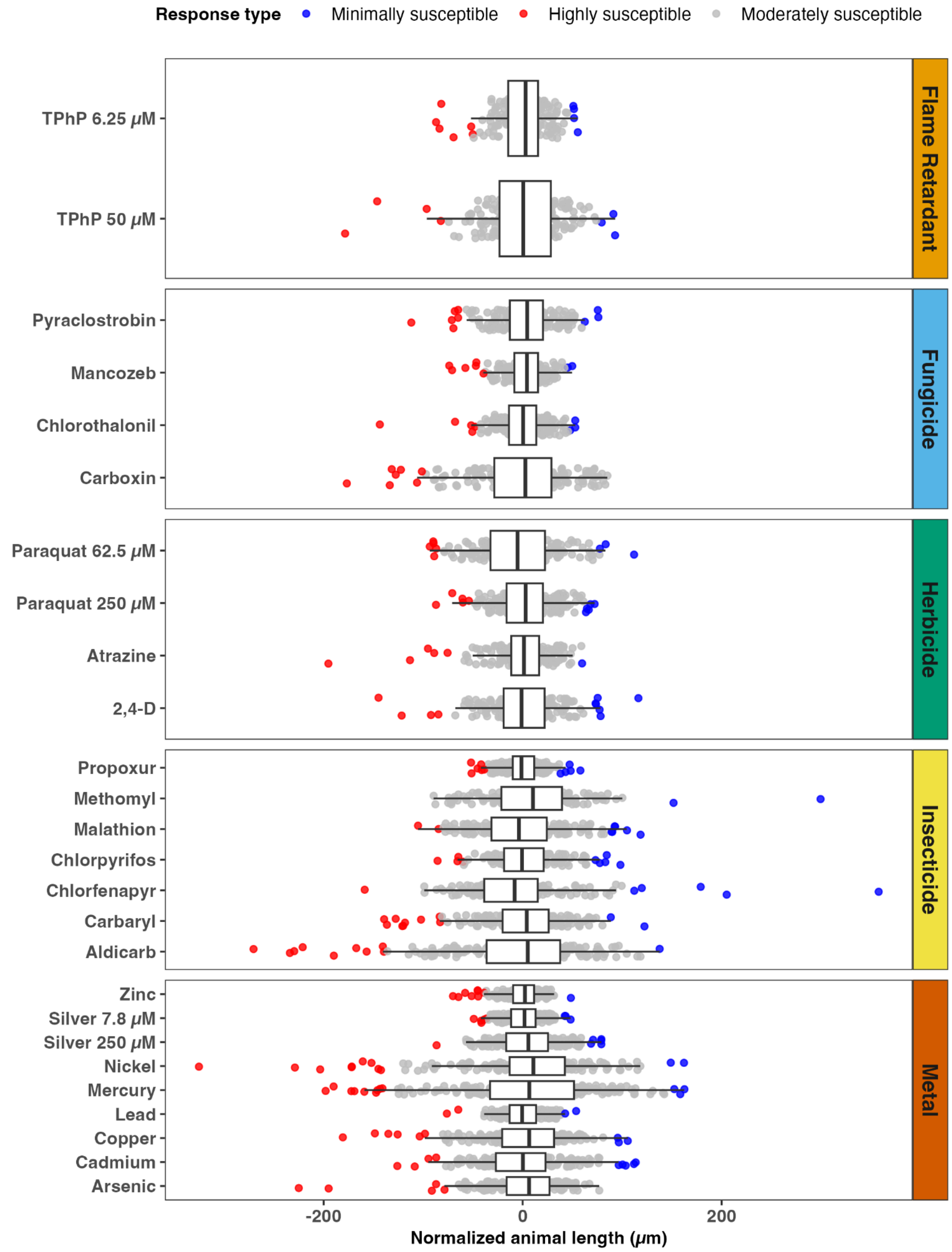


**Figure S2. Differences in susceptibility among 195 *C. elegans* strains to 26 environmental toxicant conditions.**

The normalized length differences of each strain (x-axis) in a toxicant condition (y-axis) is represented by a colored point. Toxicants are arranged along the y-axis by use group. Strains that exhibited a distinct response to a toxicant and were either highly susceptible (normalized length difference < mean -2 standard deviations) or were minimally susceptible (normalized length difference > mean +2 standard deviations) are represented by red or blue points, respectively. Strains with normalized length differences that were within two standard deviations are grey. The distribution of differences in susceptibility among all strains in each toxicant condition is shown as a boxplot. For each box, the median normalized length difference of all strains in the distribution is marked by the central line, the box edges represent the first and third quartiles, and the whiskers extend to 1.5 times the interquartile range.

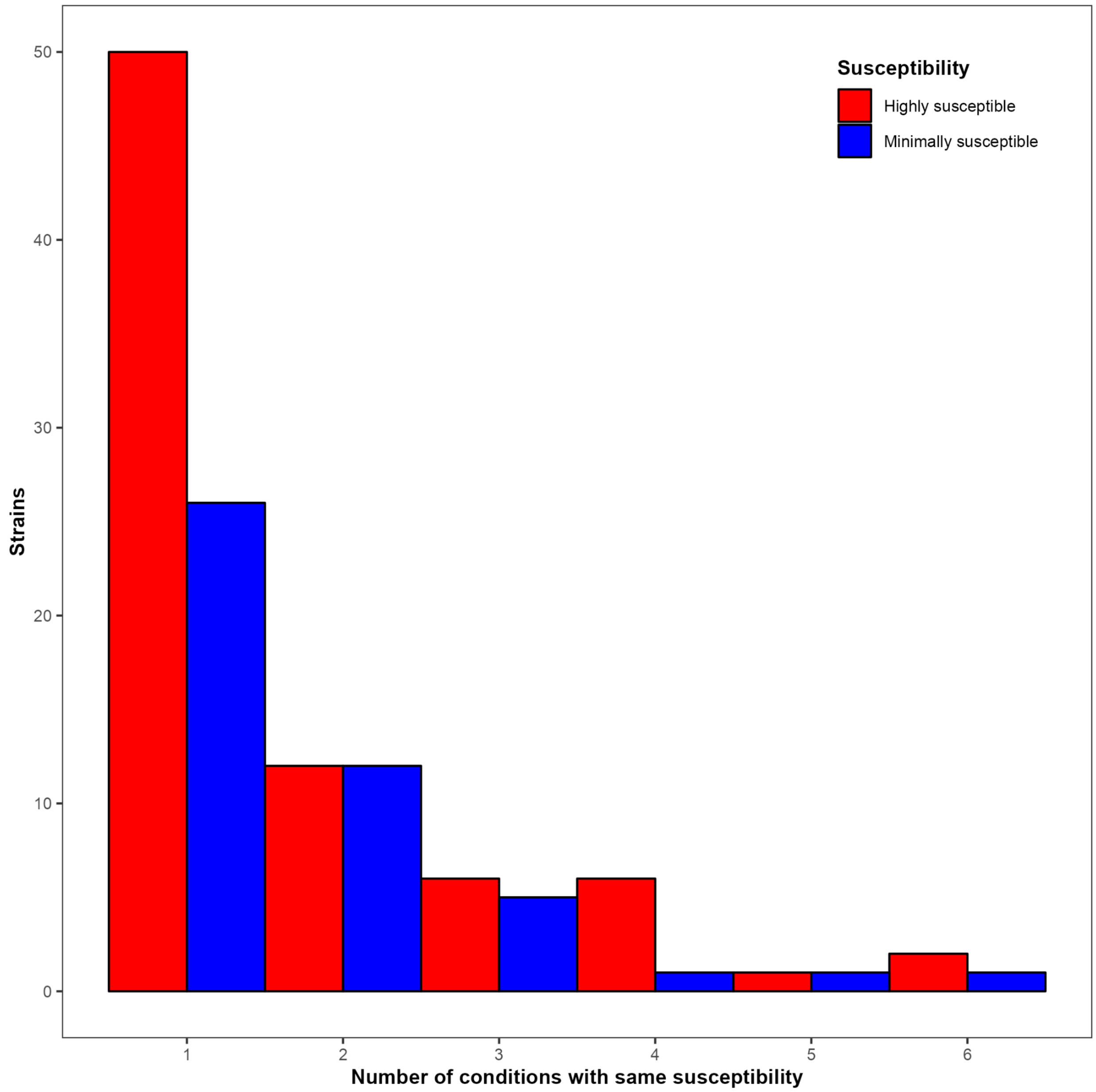


**Figure S3. Number of strains with extreme susceptibility profiles across 26 toxicant exposures.** The height of each bar represents the subset of strains (n = 108) that exhibited either highly susceptible (Red bars; normalized length difference < mean -2 standard deviations) below the mean) or were minimally susceptible (Blue bars; normalized length difference > mean +2 standard deviations) to at least one toxicant condition.

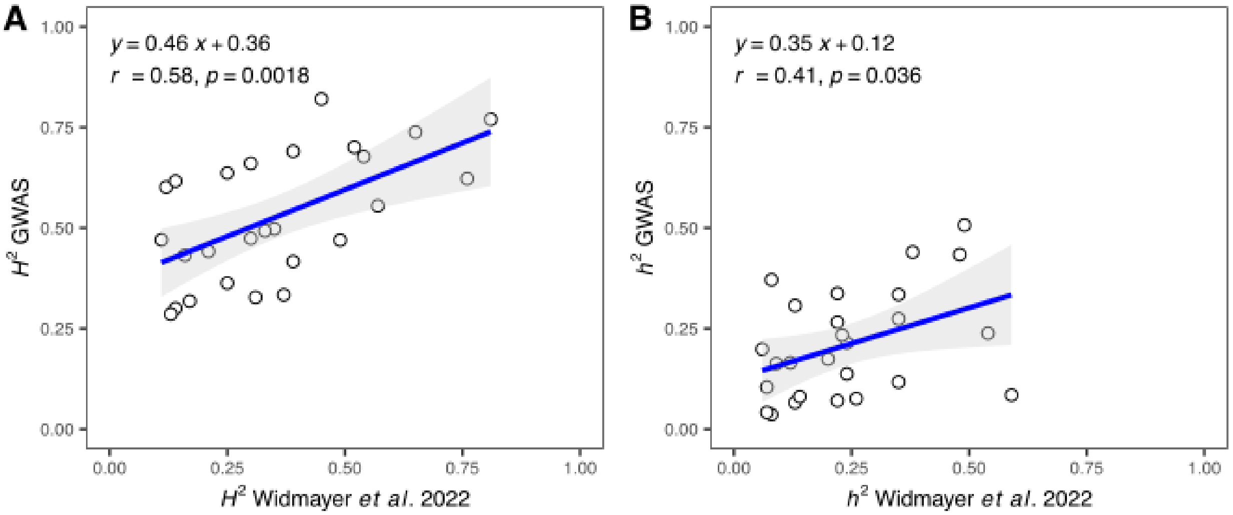


**Figure S4. Regression of heritability estimates for the 195 strains in the GWAS mapping panel against the eight strains from Widmayer *et al.* 2022.** (**A**) Broad-sense heritability estimates. (**B**) Narrow-sense heritability estimates. Each point represents the heritability estimate for a particular toxicant condition. In each panel, the regression equation, Pearson's correlation coefficient (*r*), and the associated *p*-value (*p*) are shown in the upper left corner.

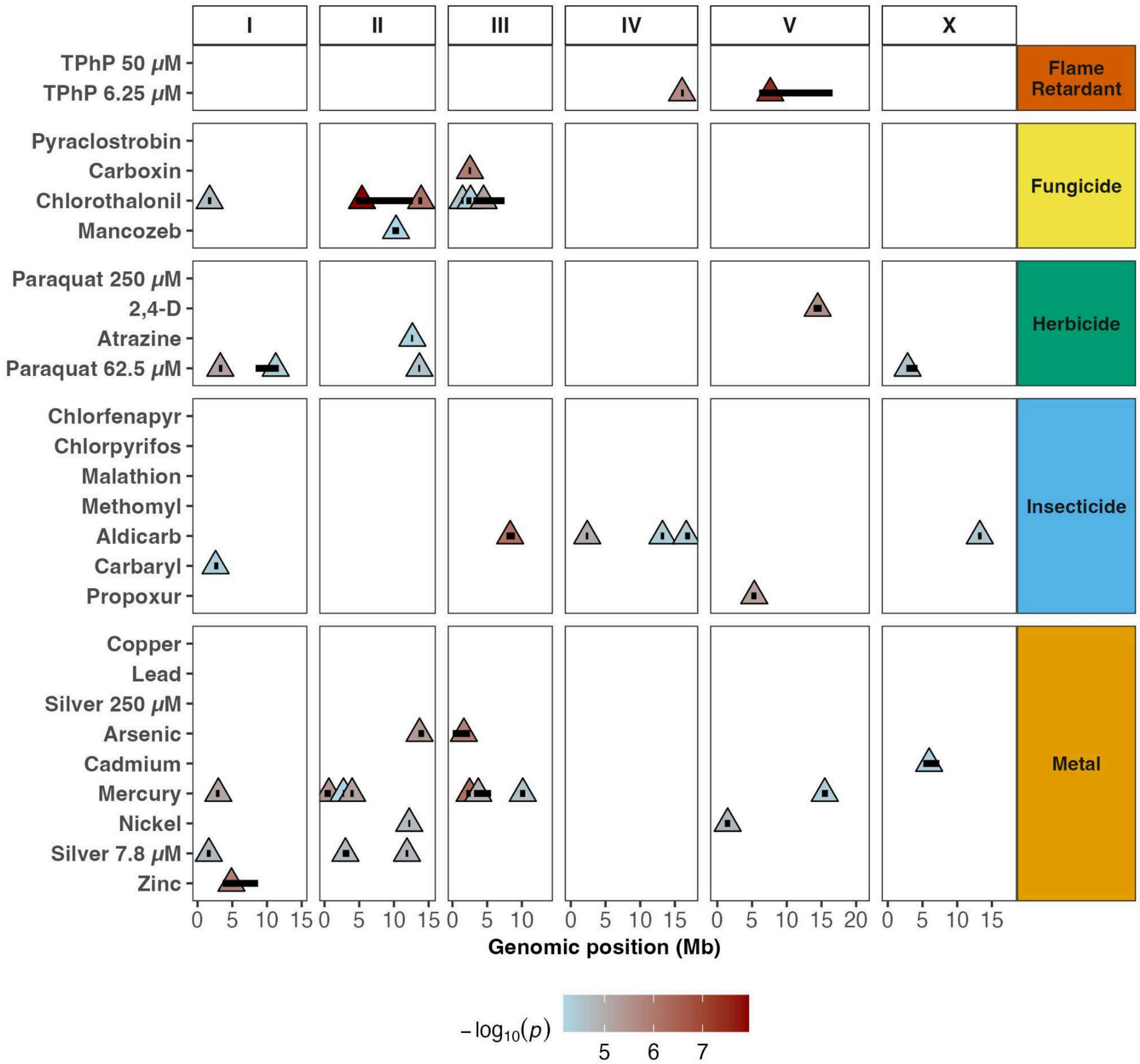


**Figure S5. Susceptibility-associated regions for all 26 toxicant exposures**. For each condition with at least one marker above the eigen significance threshold, susceptibility-associated regions are displayed by genomic position (Mb). Each region is represented by a horizontal black bar along its genomic span. The peak marker from each region is shown as a triangle colored according to its significance (-$\log_{10}$(*p*-value)). Toxicant conditions are grouped on the y-axis by use classification (Flame Retardant, Fungicide, Herbicide, Insecticide, and Metal) and displayed across the six *C. elegans* chromosomes (I, II, III, IV, V, and X) on the x-axis.

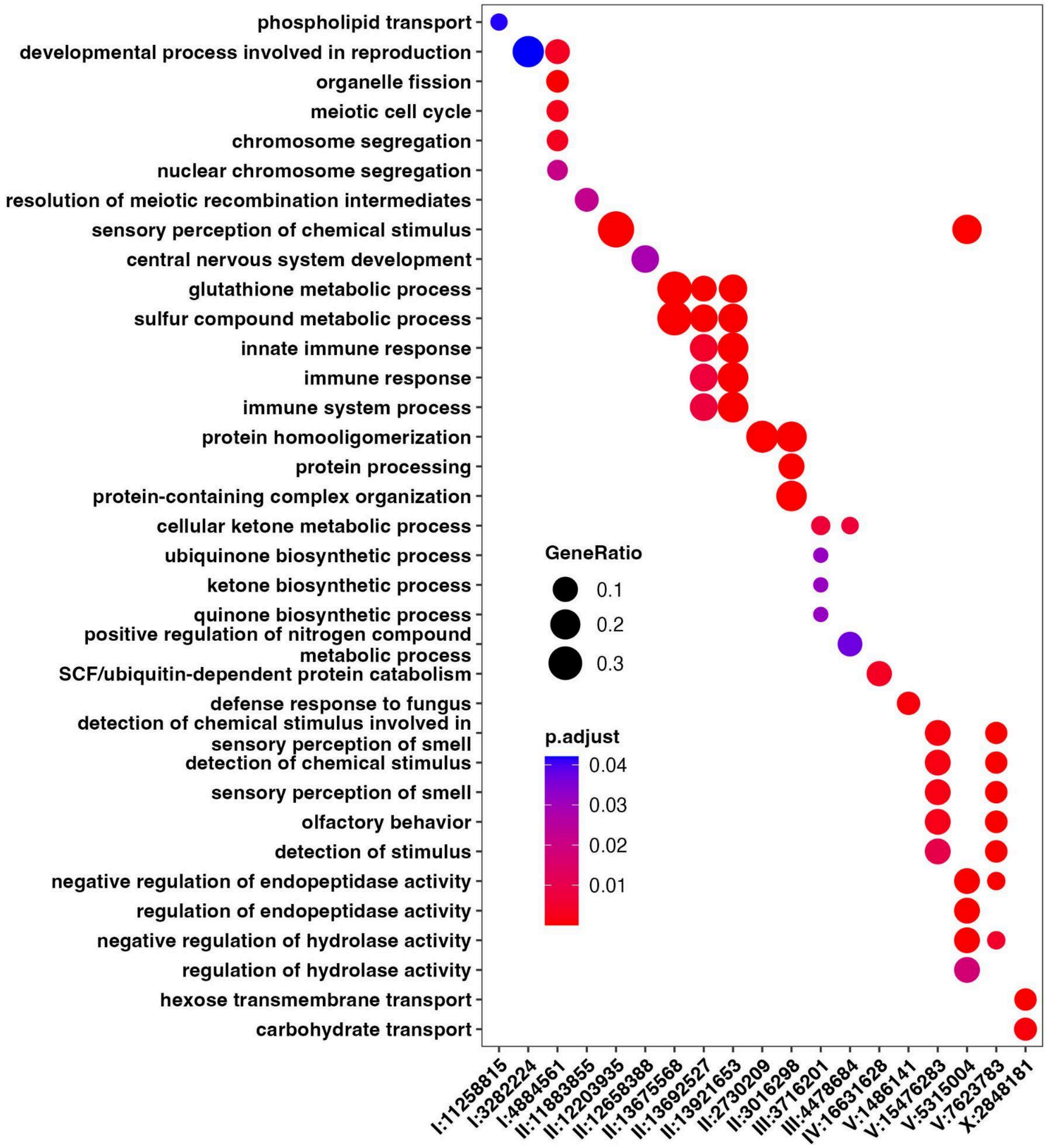


**Figure S6. Gene ontology (GO) enrichment of susceptibility-associated genomic intervals.** Biological process GO terms overrepresented by genes in susceptibility-associated intervals are visualized as a dot plot. The y-axis lists biological process GO terms that were significantly overrepresented (Bonferroni adjusted *P* value > 0.05 & *q* value > 0.05) within at least one interval. The x-axis displays the susceptibility-associated intervals, identified by their peak markers (*e.g.*, I:11258815). Only intervals where at least one biological process was significantly overrepresented are shown. Each enrichment is represented by a colored dot. The color reflects the statistical significance (Bonferroni adjusted *P* value), and the dot size represents the gene ratio (proportion of genes with the GO term in each interval). Redundant, overrepresented GO terms were reduced using the *simplify* function from the *clusterProfiler* package, which calculates semantic similarity between terms and selects one representative term when similarity exceeds 0.7. For visualization purposes, the GO term "SCF-dependent proteasomal ubiquitin-dependent protein catabolic processes" was shortened to "SCF/ubiquitin-dependent protein catabolism."

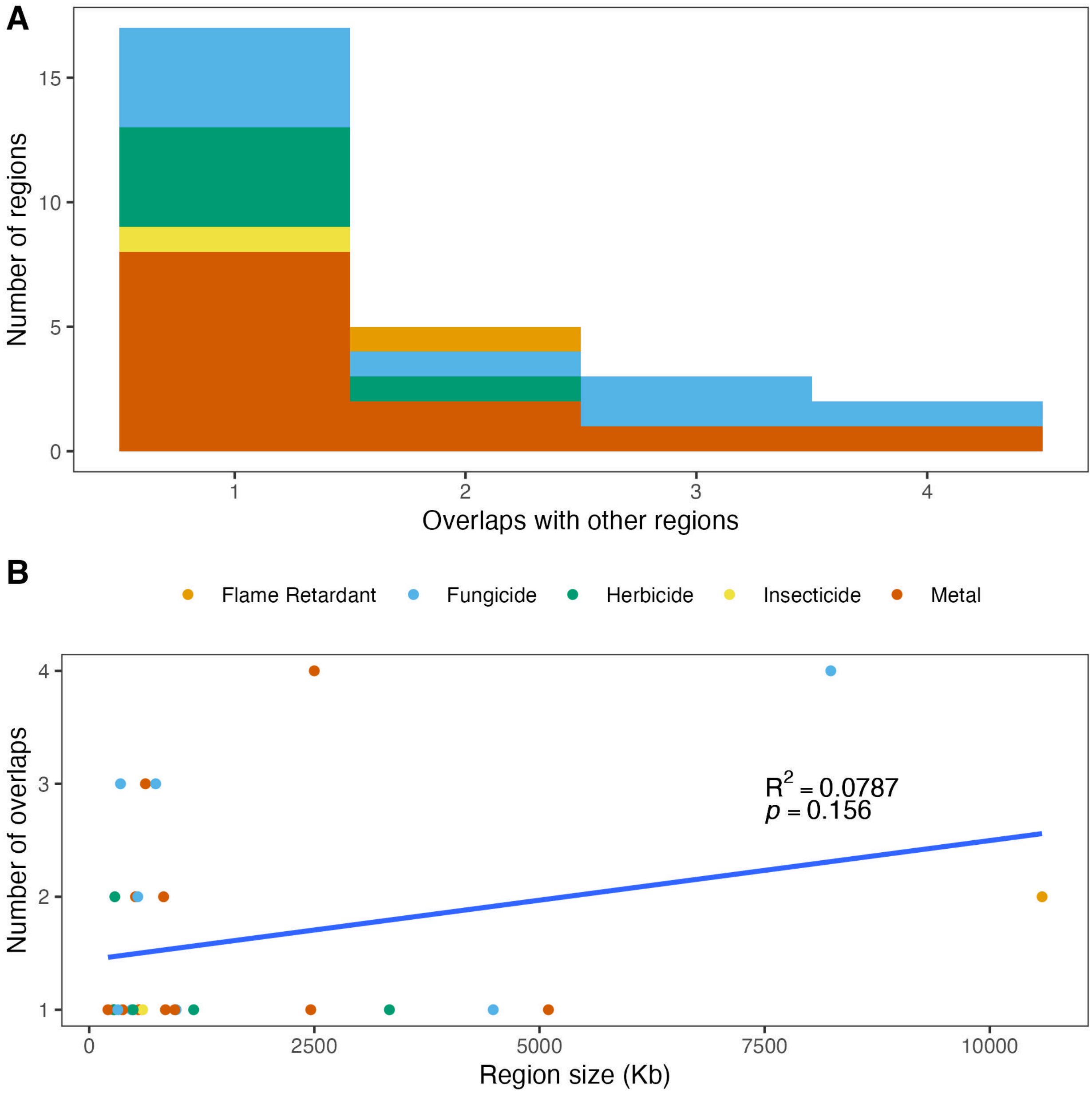


**Figure S7. Physical overlaps of susceptibility-associated intervals across toxicants**. (**A**) Histogram depicting the number of susceptibility-associated regions with which each region overlaps, limited to regions overlapping at least one other region (n = 27/40 detected regions). The bar colors indicate the toxicant class of each region. (**B**) A scatter plot that depicts the relationship between region size and the number of overlaps. Each point represents a unique region and is colored by the toxicant use group. The line shows a linear regression of the region size on the number of overlaps; the associated p-value and coefficient of determination ($R^2$) are displayed on the plot.

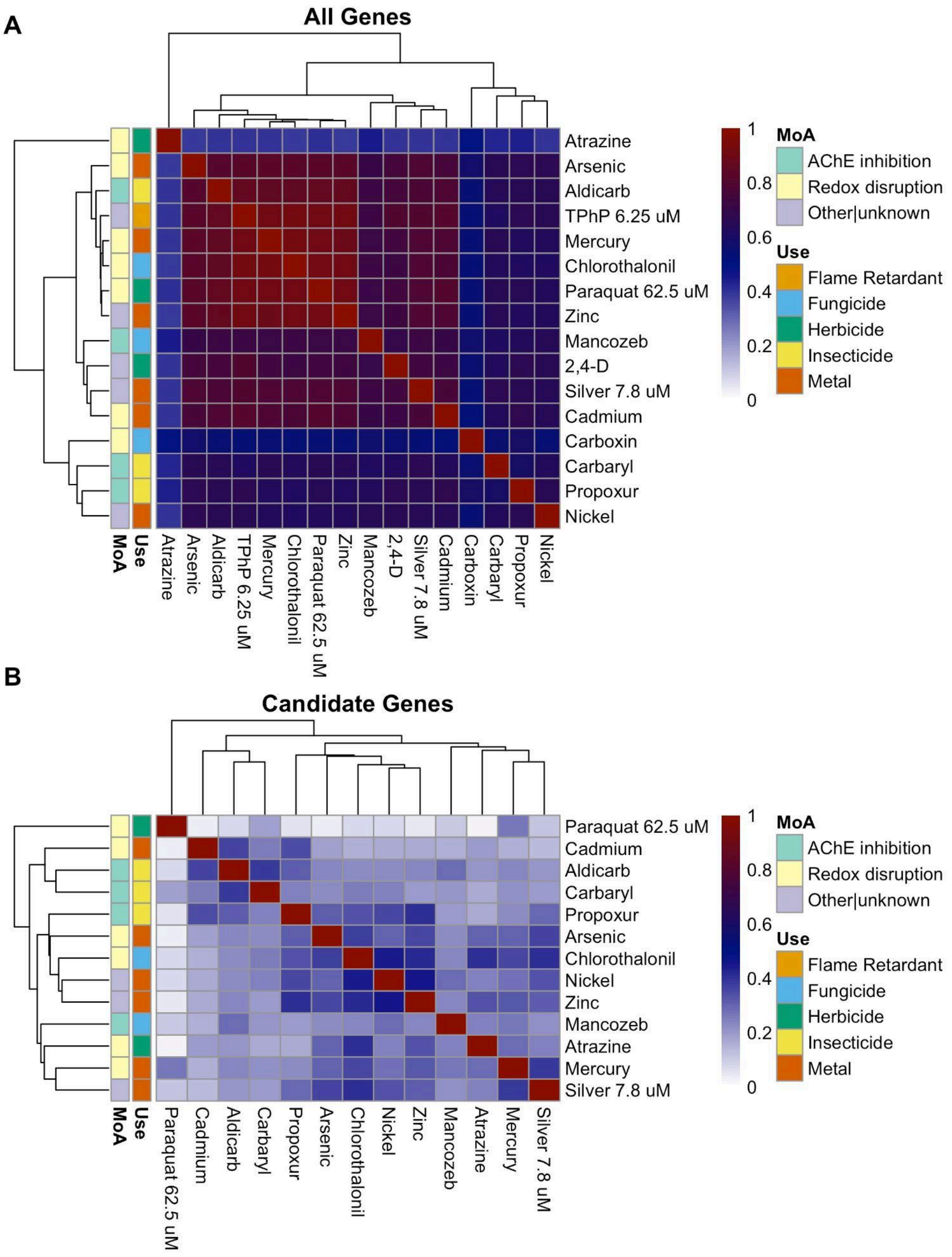


**Figure S8. GO term semantic similarity of susceptibility-associated genes.** A heatmap displays the semantic similarity of biological process GO terms for genes in susceptibility-associated intervals across multiple toxicant conditions. For each toxicant condition pair, semantic similarity was calculated using the Wang method, where higher values indicate greater functional relatedness between gene sets. Red cells indicate high semantic similarity, and blue represents low similarity. Toxicant conditions are arranged by hierarchical clustering (dendrogram displayed above the heatmap, with branch lengths that represent the similarity among toxicant conditions) so that similar toxicant conditions are displayed near each other. Vertical color-coded bars along the left side of the heatmap denote the toxicant use and MoA groups. Panel (**A**) displays the semantic similarity estimate of all genes in susceptibility-associated intervals, and panel (**B**) displays the same estimate made with only candidate genes identified for each toxicant condition.

### SUPPLEMENTAL TABLES

**Table S1. Strains selected for the GWAS mapping panel.** Alternative strain names, collection locations, and environmental parameters at collection sites are also provided where available. All strain data are taken from the CaeNDR website**.**

**Table S2. Environmental toxicants and exposure conditions assayed.** The toxicant column lists the preferred name of each toxicant retrieved by querying the CAS number against the EcoTox database. The Sigma Aldrich catalog number for each toxicant is listed along with the stock concentration prepared for each toxicant. The use group column assigns toxicants to one of five use groups. The mechanism of action (MoA) group column assigns the MoA to toxicants. Both "use" and "MoA" groups are based on a comprehensive analysis of experimental data (Kramer *et al.*, 2024). We simplified the MoA group into three categories: acetylcholinesterase (AChE) inhibition, redox disruption, and other/unknown mechanisms. Toxicants that were assayed at two concentrations are present in two rows, which distinguish the unique conditions.

**Table S3. Strain specific toxicant susceptibility data**. The relative susceptibilities of strains to each toxicant condition are provided. The strain column includes the strains included in the GWAS. The remaining columns provide the mean normalized length differences for each strain across all toxicant conditions. Missing trait data for some strain-condition combinations reflect data cleaning procedures (see methods, **Section 2.6**).

**Table S4. Differences in susceptibility to toxicant conditions among strains.** The table displays the standard deviation (SD) of the normalized length differences. Highly susceptible strains (normalized length difference < mean -2 standard deviations) and minimally susceptible strains (normalized length difference > mean +2 standard deviations) are listed.

**Table S5. Comparison of the mean standard deviation of normalized length differences across toxicant groups**

**Table S6. Heritability estimates for toxicant conditions.** For each toxicant condition, broad-sense heritability ($H^2$) and narrow-sense heritability ($h^2$) estimates are presented with their 95% confidence intervals (CI) [Upper CI, Lower CI] generated with bootstrapping.

**Table S7. Susceptibility-associated intervals detected by the Inbred GWAS algorithm across toxicant conditions.** The table presents genomic intervals associated with natural differences in toxicant susceptibility identified by genome-wide association mapping with the Inbred algorithm. Each interval is represented by its most significant marker (Peak Marker) and defined by its genomic coordinates (Interval). Intervals are grouped by toxicant condition and ordered by statistical significance. For each interval, the statistical significance (-log10(p-value), the effect size, which represents the change in normalized length difference between strains with alternative alleles, the frequency of the alternative peak marker in the 195 strains, and the proportion of differences in susceptibility explained by the marker.

**Table S8. Candidate genes from susceptibility-associated intervals detected by GWAS.** The table lists the 136 candidate susceptibility genes from genomic intervals identified by GWAS with the Inbred algorithm. For each candidate gene, the table displays the strength of the association (-log10(*p*-value) between a high-impact single-nucleotide variant (SNV) in each gene with differences in toxicant susceptibility, the linkage disequilibrium ($r^2$) of that SNV with the peak marker, and the frequency of that SNV in the strains assayed. The interval column indicates the toxicant condition and the specific susceptibility-associated interval where each candidate gene was identified.

**Table S9. Overlapping susceptibility-associated intervals across toxicant condition pairs.** The table displays toxicant condition pairs that share at least one physically overlapping susceptibility-associated interval. For each pair, the overlapping intervals are identified by their peak marker and the linkage disequilibrium ($r^2$) estimate between peak markers for each pair of overlapping intervals.

**Table S10. Prioritization of actionable susceptibility-associated regions detected by GWAS.** The 22 susceptibility-associated genomic regions with at least 20 strains carrying the alternative peak marker allele, ranked from most actionable (1) to least actionable (22) according to a hierarchical prioritization scheme. Regions were first ranked by statistical significance (-log10(*p*-value)). Subsequently, regions were categorized based on their linkage disequilibrium (LD) structure with manual curation of fine-mapping results (<250kb, 200-500Kb, or >500kb). Regions with lower LD structure (<250kb) classifications were prioritized for their higher mapping resolution compared to those with more extensive LD structure (>500kb). Finally, regions were prioritized based on the proportion of strains with hyper-divergent regions (HDRs) that fell within or overlapped the interval. A lower proportion of strains with the alternative peak marker allele with HDR overlap resulted in higher prioritization. For each interval, the table displays the toxicant condition and peak marker position, statistical significance, LD structure, reference and alternative peak marker allele frequencies, and the proportion of strains with the alternative allele with at least one HDR that overlapped the interval.

**Table S11. Human orthologs for candidate genes.** The 94 actionable candidate genes are listed as in Table S7. Human orthologs identified by one or more orthology-prediction programs used in ortholist2 are shown in the column titled “HGNC Symbol”. All descriptive columns from ortholist2 data are also included.

### SUPPLEMENTAL METHODS

#### Methods S1. Detailed image processing and data cleaning steps

We first removed nematode objects with a length less than 100 µm because these are typically not nematodes as described in (Widmayer *et al.*, 2022; Shaver *et al.*, 2023). Second, used the *modelSelection*, *edgeFlags*, and *setFlags* functions with default parameters from *R/easyXpress* (v1.0.0) (Nyaanga *et al.*, 2021) to filter out nematode objects near the well edge and drop nematode objects if they occur in the same ROI. These filters reduce the number of improperly segmented animals that occur from illumination irregularities near the well edge and the number of spurious nematode objects identified in ROIs that contain debris, illumination artifacts, and/or nematode clusters. Third, we increased the minimum length threshold from 100 µm to 165 µm for experimental conditions in which this adjustment reduced the fraction of spurious small objects. To determine if the higher length threshold improved data quality, we manually reviewed putative nematode objects detected under both thresholds (100 µm and 165 µm) for the four strains with the smallest mean animal length in each condition.

Despite applying the three filtering steps above, we noticed spurious objects persisted in the data. Because manually filtering these anomalies would require reviewing millions of objects, we sampled a total of 4,500 objects across nine different axes of variation (object integrated intensity, object minimum edge intensity, object mean edge intensity, ROI compactness, ROI form factor, CellProfiler model type selected, number of objects detected in image, image illumination consistency, variation in object number detected within condition), in our data and manually scored these objects as “nematode” or “non-nematode”. The “nematode” class describes a ROI that was a single nematode and had a single CellProfiler model fit within the ROI that covered at least 90% of the animal’s true length. All other cases were classified as a “non-nematode”. The types of spurious “non-nematode” data included larger debris, clumps of smaller nematodes, nematodes that were not properly segmented from the image background, and CellProfiler models that simply did not fit the ROI properly (*i.e.*, less than 90% of the animal's true length). To sample the 4,500 objects from our full data set, we wrote the *makeTrainingSet* function. This function produces a reproducible sample of objects across whatever input variables are supplied when the seed parameter is set, *e.g.*, seed=100. To visualize the objects and assist in manual scoring, we input those objects into the *viewObjects* function to output i) individual object images that can be scored manually and ii) a comma-separated value file with object IDs to assist in tracking the manual scoring. Next, 30% of the manually scored data was used to train a stochastic gradient boosted classifier with the *caret* R package (v6.0-92) (Kuhn, 2008). The resulting classifier model was saved and subsequently used to filter out "non-nematode" objects in our data. The classifier accuracy was 89.6%, which was estimated by resampling the training data with 10-fold cross-validation.

After the classifier filter, we removed nematode objects with outlier lengths within each well. The outliers were detected using Tukey's fences (Tukey, 1977). We then calculated summary statistics for each well using the filtered nematode objects. The summary statistics included the total number of nematode objects in each well, median, mean, standard deviation, and the coefficient of variation (CV) of the nematode object lengths.

Next, we removed data from a particular bleach synchronization of a strain if we found a high degree of variation in the number of nematode objects within control wells for that bleach synchronization. We measured the variance by grouping the data by bleach and strain, then calculating the CV in nematode object counts across all the control wells. This filter was used because we reasoned that unusual variance in animal densities among replicate wells could have occurred from improper bleaching or embryo titration, and could potentially bias our estimates of toxicant responses. The exact CV threshold for removing wells (CV > 0.68) was determined by inspecting the distribution of CV values and selecting a threshold that excluded only the extreme values. We then filtered out data from wells with fewer than five, or greater than 30 nematode objects that passed our previous filters. On the low end, this filter ensured that we had enough nematode objects within the well to get a good estimate of well summary statistics. On the high end, this filter helped us remove wells with high animal density or possible contamination, which could bias our estimate of toxicant responses. Afterwards, we grouped the data by strain and condition and removed wells with outlier estimates of median animal length using Tukey's fences.

Next, we normalized our data by subtracting the median animal length in control conditions for each strain in each assay and bleach from the median animal length in the corresponding treatment wells for the same strain. We refer to those normalized values as "control delta" traits. Strains have natural differences in growth in control conditions, and the control delta normalization allowed us to remove the effect of those strain differences and retain the effects of assay and bleach in the data. Next, for each condition, we estimated the effect sizes of each assay and bleach by extracting the regression coefficients from a linear model of the formula *control_delta ~ assay_bleach - 1*. We then plotted the effect sizes to observe outlier assays and bleaches with extreme effects. We filtered out all data from assays and bleaches with extreme effect sizes within a condition. We then corrected for any remaining assay and bleach effects within a condition by taking the residual values from the linear model of the formula *control_delta ~ assay_bleach - 1*. Finally, we removed all data for a strain in a particular condition if we retained data from fewer than three wells after applying all the filters above.

**Methods S2. Identification of hyper-divergent regions (HDRs) in *C. elegans***

Hyper-divergent regions (HDRs) were identified using methods described previously (Lee *et al.*, 2021) with modifications. First, we aligned the 15 wild strain genomes assembled from long-read data (Lee *et al.*, 2021) against the N2 reference genome (WormBase, WS283) (Sternberg *et al.*, 2024) using *nucmer* v3.1 (custom parameters --mingap 500 --mincluster 100) (Kurtz *et al.*, 2004). The N2 reference genome was then partitioned into 1 kb bins using *bedtools* v2.31.1 *makewindows (Quinlan and Hall, 2010)*. For each strain, the identity of every bin was estimated from the average identity of the alignment spanning the bin. When multiple alignments of variable length were mapped to a single bin, the alignment with the longest span was used to estimate the identity. To identify hyper-divergent (HD) bins, instead of the coverage fraction (proportion of read depth of the bin relative to the genome-wide depth average) threshold used by *Lee et al*. we used the percentage of bases covered by at least one aligned read (1x coverage) in each bin (percent bases covered). Using the previous identity threshold, bins under 95% identity were classified as hyper-divergent (Lee *et al.*, 2021). Bins with less than 60% bases covered were also classified as HD. An overwhelming majority of bins had 100% or 0% bases covered, and the 60% bases covered threshold simply polarized bins to identify gaps in coverage with a boundary uncertainty of 1 kb (bin size). To further refine HDR boundaries, non-HD bins that were flanked by two HD bins were re-classified as HD. Contiguous bins classified as HD were clustered into HDRs. HDRs separated by less than 5 kb were merged, and HDRs smaller than 5 kb were filtered out. HDRs that were clustered exclusively from bins under 60% bases covered were also filtered out to prevent deletions from being included among HDRs.

Next, to determine the optimal parameters to call HDRs species-wide using short-read sequencing data, we first aligned the Illumina sequences of 611 wild strains (CaeNDR release 20231213) against the reference genome, and called variants using GATK (van der Auwera and O'Connor, 2020). For each strain, we estimated the SNV count using *bcftools* v1.21 (Danecek *et* *al.*, 2021) and *bedtools coverage* v2.31.1 (Quinlan and Hall, 2010), and the percentage of bases covered in every reference bin using mosdepth v0.3.10 (Pedersen and Quinlan, 2018). We tested a wide range of variant counts (5 to 25 SNVs per kb, 1 SNV step) and percent bases covered (5 to 90%, 5% step) thresholds to classify and cluster bins into HDRs using short-read data for the 15 strains where we previously generated long-read HDR calls (Lee *et al.*, 2021). We selected the optimal variant count and base coverage thresholds for short-read HDR calling by estimating the concordance between short- and long-read HDR calls. We estimated the concordance of short- and long-read HDR calls by measuring the overlap fraction (proportion of a long-read HDRs that overlapped a short-read HDR) and the excess fraction (proportion of a short-read calls that exceeded the boundary of an overlapping long-read call). For each strain, we estimated the mean overlap fraction and mean excess fraction across every overlap between the short- and long-read HDR calls at every threshold pair. We also estimated the completeness and accuracy of the short-read HD calls by calculating recall (proportion of long-read calls that had any overlap with short-read calls) and precision (proportion of short-read calls that had any overlap with long-read calls). We selected the top N (starting at N=1) threshold pairs based on the mean overlap fraction for each strain, and increased the value of N by one until we identified a threshold pair that was present among all of the 15 strains in this comparison (referred to as 'consensus optimal', reached at N=40, top 11% of all threshold pairs). Threshold pairs with a variant count threshold lower than eight SNVs were excluded from the search of a consensus optimal because of substantially lower precision and increased mean excess fraction, at the expense of small reductions to recall and mean overlap fraction. The consensus optimal thresholds selected were a variant count of 11 SNVs per kb paired with a percent bases covered of 80%. We classified bins with estimated variant count or percent bases covered under these optimal thresholds as HD, genome-wide, across all 611 wild strains. We then performed the same bin clustering, gap re-classification, region clustering, and size and coverage filtering steps that were applied to regions called with long-read sequencing data.
